# Pharmacological and genetic access to astrocytic cholesterol programs in human neural cells

**DOI:** 10.64898/2026.11.16.767064

**Authors:** Matthew Tegtmeyer, Noah Pettinari, Ajay Nadig, Jim Nemesh, He Jax Xu, Vy Ngo, Marina Hogan, Sonia Bolshakova, Dhara Liyanage, Emily Peirent, Adrianna Maglieri, Derek Hawes, Kiku Ichihara, Steven A. McCarroll, Ralda Nehme

## Abstract

Astrocyte–neuron interactions coordinate cholesterol metabolism, cell adhesion, and synaptic gene expression ^1,2^, and related gene expression programs decline together in schizophrenia^3^. Whether pharmacological or genetic perturbations can selectively engage components of this coordinated biology was unknown. Here we combine genetically diverse cultures of human stem-cell-derived neurons and astrocytes, pharmacological profiling, genetic association mapping and CRISPR interference to examine these biological programs. In stem-cell-derived astrocyte mono-cultures, the antipsychotic clozapine induced a focused transcriptional response involving cholesterol biosynthesis and export, and drove an increase in intracellular and secreted cholesterol. This response overlapped with the cholesterol-related component of the astrocytic response to neuronal contact seen in neuron-astrocyte co-cultures, but did not extend to synaptic cell-adhesion genes. In neuron–astrocyte co-cultures, clozapine also increased neuronal firing. Across individual cell donors, cholesterol-pathway induction by clozapine was broadly shared but varied in magnitude. Gene-expression patterns in untreated donors were associated with differential responses to clozapine and the liver X receptor agonist GW3965. Common genetic variation influenced expression of several cholesterol-related genes, including a clozapine-dependent association of *HSD17B7* expression with nearby genetic variants. Independently, repression of *LDLR* induced coordinated expression of cholesterol-biosynthesis genes in astrocytes that overlapped with astrocyte responses to clozapine and neuronal contact. These findings identify shared cholesterol-related transcriptional responses to pharmacological and genetic perturbations and link common genetic variation to distinct features of cholesterol-pathway responses.

## Introduction

Schizophrenia has a complex genetic architecture, with common and rare variants implicating genes involved in synaptic organization, neuronal differentiation and neurotransmission ^4,5^. These findings have identified biological processes relevant to illness, but how those processes relate to the cellular effects of antipsychotic treatment remains unclear. Central questions include: whether pharmacological treatments engage coordinated cellular programs implicated by schizophrenia genetics and post-mortem studies; which components of those programs respond to treatment; and whether these cellular responses vary across individuals.

Astrocyte–neuron interactions provide a context in which to examine this relationship. Astrocytes support synaptic development through both physical interactions with, and the provision of metabolites like cholesterol to neurons. Experiments in neuron-astrocyte co-cultures identified cholesterol carried by apolipoprotein E-containing lipoproteins as an astrocytic signal that promotes the formation of functional synapses^6^. Astrocyte-secreted thrombospondins and glypicans also promote distinct aspects of excitatory synapse formation ^7,8^. *ABCA1* contributes to this lipid-supply system by supporting the lipidation of astrocyte-secreted *APOE* ^9^. Coordinated astrocytic and neuronal expression is also evident in the human brain. Single-nucleus RNA-seq analysis of human prefrontal cortex identified a synaptic neuron and astrocyte program (SNAP), in which neuronal expression of synaptic genes covaries across individuals with astrocytic expression of genes involved in synaptic adhesion, neurotransmitter handling and cholesterol biosynthesis ^3^. Both the neuronal and astrocytic gene-expression programs in SNAP recruit sets of genes enriched for schizophrenia genetic associations and show reduced expression in schizophrenia and with advanced age. These observations connect astrocyte–neuron coordination to disease-associated biology, while inviting the question of how its constituent processes can be modulated genetically and pharmacologically.

Co-culture experimental systems provide access to these coordinated responses. In human iPSC-derived neurons, contact with mouse astrocytes induced neuronal synaptic expression programs, which were associated with astrocytic expression of cell-adhesion genes, including genes implicated in schizophrenia ^1^. Time-resolved profiling shows that induction of astrocytic cholesterol-biosynthesis genes precedes activation of neuronal synaptic programs, and that repressing neuronal cholesterol-uptake receptors reduces synaptic gene expression and alters the accompanying astrocyte response^2^. Together, these studies identify cholesterol metabolism and synaptic adhesion as separable (though sometimes coordinated) components of astrocyte–neuron interactions. Whether pharmacological perturbations preferentially engage one component, and how their effects correspond to the broader response to neuronal contact, are unknown.

A further question is how these cellular responses vary across individuals. Common genetic variation contributes to molecular heterogeneity among human iPSC lines ^10^. A shared pathway response may differ in magnitude according to baseline transcriptional state or genetic background. Pooled cultures of cells from multiple donors, or cell villages, allow donor-specific expression and phenotypes to be measured under shared experimental conditions, providing a means to examine natural variation in baseline state and perturbation responses ^11,12^. Complementary pooled CRISPR screens with single-cell transcriptomic readouts enable direct assessment of the expression changes that follow targeted genetic perturbation ^13,14^. Combining these approaches allows pharmacological responses, common regulatory variation and targeted gene repression to be examined within related experimental systems.

Here we combine multi-donor human iPSC-derived neural cultures, pharmacological perturbation, cis-eQTL mapping and CRISPRi to examine how schizophrenia-relevant perturbations engage astrocyte–neuron expression programs. The antipsychotic clozapine preferentially induced the cholesterol-related component of the astrocyte response to neuronal contact, accompanied by increased intracellular and conditioned-medium cholesterol. Across donors, cholesterol-pathway induction was broadly shared but varied in magnitude, and baseline gene-expression patterns were associated with differential responses to clozapine and the LXR agonist GW3965. Common genetic variation influenced expression of several cholesterol-related genes, with a treatment-dependent association at *HSD17B7*. In astrocytes, *LDLR* repression induced coordinated cholesterol-biosynthesis transcription overlapping the responses to clozapine and neuronal contact, while neuronal perturbations produced distinct changes in synaptic and translation-related programs. These findings define experimentally accessible components of astrocyte–neuron interactions and reveal how baseline expression and genetic variation relate to their responses to perturbation.

### A perturbation screen across donors, treatments, and cell types

We optimized and used cell villages, in which iPSCs from multiple donors are pooled, differentiated, and profiled together. Computational methods such as Census-seq and Dropulation then measure donor representation from DNA or RNA sequencing, respectively, and assign individual cells to donors in single-cell RNA sequencing data ^11,15^. Lines carrying known growth-promoting variants^16^ or karyotypic abnormalities were excluded before pooling. We analysed donor representation across three independent village experiments, and found that donor proportions changed during extended culture and passaging of iPSC villages, consistent with previous findings^15^. However, donor proportions remained correlated and largely unchanged before and after cryopreservation, enabling the banking of multiple iPSC village vials for use in downstream experiments. When mixed at the iPSC stage, cells differentiated into neuronal progenitor cells ^17^ (NPC, day 4 post-neuronal induction) or into fully differentiated neurons ^17^ (day 28 post-neuronal induction) retained mostly balanced donor representation, whereas the astrocyte^18^ samples showed greater donor imbalance after extended differentiation (day 30 post-induction), during which astrocytes, unlike neurons, continue to proliferate (Extended Data Figure 1; Supplementary Table 1). Overall, our optimized workflow enabled the donor-resolved screen we present below. Our results indicate that while iPSC villages with large numbers of donors are tractable for endpoint differentiated cell types that pass through only a limited proliferation stage, they require additional optimization for cell types that proliferate for long periods before reaching the required differentiated state, such as astrocytes, in line with our previous findings in brain organoid villages, or chimeroids^19^.

To ask which pharmacological perturbations engage astrocyte–neuron biology, we profiled transcriptional responses to a panel of seventeen perturbation conditions spanning synaptic signalling (glutamate, GluN2A PAM), calcium homeostasis (isradipine), antipsychotic pharmacology (clozapine, haloperidol), oxidative stress (hydrogen peroxide), antiviral exposure (azidothymidine, efavirenz), inflammatory signalling (TNF-*α*, IFN-*α*, IFN-*γ*, IL-1*α*, C1q) and lipid metabolism (atorvastatin, simvastatin), in pooled multi-donor iPSC-derived neurons (cultured with mouse astrocytes from D4 to promote maturation) and in multi-donor iPSC-derived monoculture astrocytes from 44 genetically distinct donors (Figure 1a; Supplementary Tables 2 and 3). Single-cell RNA sequencing across more than 2.3 million cells was the primary readout. The screen cohort included 28 male and 16 female donors with a median age of 60 years and a range of 21-93 (Extended Data Figure 1a-c; Supplementary Table 1). The donor-representation analyses and their experimental scope are described in Methods and Extended Data Figure 1d-h. To understand how neurons and astrocytes responded to these treatments, we asked three main questions: how similar, or different, was the overall impact of these perturbations, how much of that impact was cell-type-dependent, and how consistently did a given perturbation act across genetically distinct individuals.

**Figure 1.**
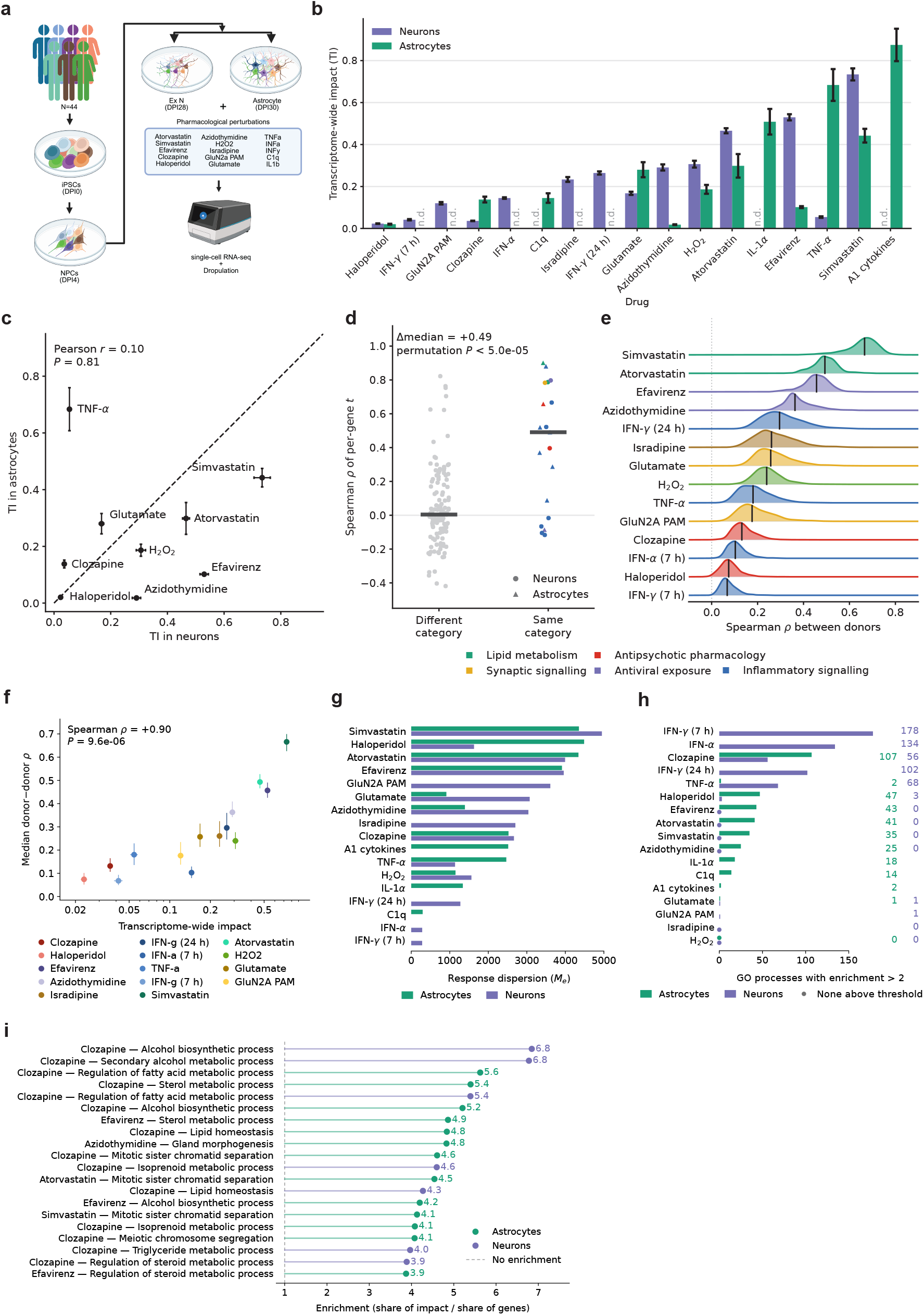
A multi-donor perturbation screen resolves treatments by functional category. **(a)** Screen schematic. A 44-donor iPSC village was differentiated in pools into neurons and astrocytes, exposed to 17 pharmacological, inflammatory and metabolic perturbation conditions, and profiled by single-cell RNA sequencing. Donors were assigned using Dropulation. **(b)** Transcriptome-wide impact (TI), estimated using TRADE, for each condition and cell type. Bars show estimates *±*1 s.e.; n.d., not determined. **(c)** Astrocytic versus neuronal TI for nine perturbations profiled in both cell types (Pearson *r* = 0.10, *P* = 0.81). Error bars indicate *±*1 s.e.; dashed line denotes equality. **(d)** Pairwise Spearman correlations of gene-level *t* statistics between perturbations within each cell type. Same-category pairs were more similar than different-category pairs (median *ρ* = 0.491 versus 0.004; permutation *P ≈* 5 *×* 10*^−^*^5^). Colors indicate functional category, shapes indicate cell type, and horizontal bars show medians. **(e)** Distributions of between-donor correlations in neuronal response profiles across 34 donors (561 pairs per perturbation), ordered by median agreement. Colors indicate functional category; black lines mark medians. **(f)** Median between-donor agreement versus TI for 14 neuronal perturbations (Spearman *ρ* = 0.90, *P* = 9.6 *×* 10*^−^*^6^). Horizontal error bars indicate *±*1 s.e. for TI; vertical bars indicate the interquartile range of donor-pair correlations. **(g)** Effective number of differentially expressed genes (*M* _e_) for each condition and cell type, summarizing response dispersion. Values range from 283 to 4,948. **(h)** Number of Gene Ontology biological-process gene sets with greater than twofold TI enrichment among 1,571 sets tested. Counts appear at right; dots indicate zero qualifying sets and blank slots indicate unprofiled conditions. **(i)** Twenty gene-set combinations with the highest TI enrichment across eight pharmacological conditions, colored by cell type. Enrichment is the fraction of TI attributed to a set divided by its fraction of genes; dashed line marks 1. Fifteen combinations involve sterol, lipid, fatty-acid or isoprenoid terms, and fourteen involve clozapine.

Perturbations differed by roughly fifty-fold in transcriptome-wide impact (TI), measured using TRADE^20^ (0.018–0.875 across 26 condition-by-cell-type estimates; Figure 1b; Supplementary Table 4). Among nine conditions measured in both cell types, TI showed little cross-cell-type correlation (Figure 1c; Pearson r = 0.096, P = 0.81). For example, TNF-*α* had a larger impact in astrocytes than neurons (TI = 0.68 versus TI = 0.05), whereas efavirenz had a larger impact in neurons (TI = 0.53 versus TI = 0.10).

The perturbations also varied in which genes they altered, and drugs from similar classes impacted similar biological programs. When we examined the gene-level correlation of their effect estimates, compounds of the same category were more similar to one another than those of a different class (within a cell type, median *ρ* = +0.491 across 19 same-class pairs against +0.004 across 125 different-class pairs; compound-category permutation P = 5 *×* 10*^−^*^5^; Figure 1d). Clozapine’s closest neighbor was haloperidol in both cell types (*ρ* = 0.658 in astrocytes, 0.396 in neurons), and the two statins were the tightest pair amongst the panel (*ρ* = 0.901 in astrocytes).

Donors varied in their responses to each perturbation. For each perturbation, we correlated gene-level response profiles between each pair of donors. Median between-donor agreement ranged almost ten-fold across the fourteen perturbations, from *ρ* = 0.068 for IFN-*γ* at 7 h to *ρ* = 0.666 for simvastatin (Figure 1e). Between-donor agreement was greater for perturbations with larger transcriptome-wide effects. We observed strong correlations between the donor spearman correlations and the magnitude of effect on the transcriptome (Figure 1f; *ρ* = +0.90, *P* = 9.6 *×* 10*^−^*^6^).

The TI measures how much a perturbation alters genome-wide gene expression, but not how widely that effect is distributed. When we measured the effective number of differentially expressed genes (Me) from our TRADE output, we observed a 17.5 fold difference across the panel of perturbations (Figure 1g). In neurons, for example, IFN-y at 7h affected a small number of genes (Me = 283) while simvastatin moved many more (Me = 4,948). Response dispersion showed little association with TI (*ρ* = 0.26, P = 0.20). Given that the TI is distributed broadly across the transcriptome, we asked how much of each perturbation’s response was concentrated into defined biological programs. For each perturbation, we computed the fraction of TI attributed to each GO biological-process set divided by the fraction of tested genes belonging to that set (Methods). We observed that some perturbations induced gene expression responses which did not cleanly map onto specific biological programs while others, such as IFN-*γ* and IFN-*α*, had many concentrated effects (Figure 1h).

We next examined what biological programs the perturbation effects were enriched for. Restricting our analysis to the eight pharmacological perturbations, excluding the IFN-*γ* and IFN-*α* treatments which we describe in detail elsewhere^21^, clozapine accounted for 14 of the top 20 most concentrated drug-by-cell-type-by-program combinations, and was heavily enriched for sterol and lipid metabolic programs in both neurons and astrocytes (Figure 1i). Efavirenz also accounted for three of the twenty highest-ranked combinations, involving sterol metabolism and alcohol biosynthesis in astrocytes (Figure 1i).

Together, these data show that perturbations differ substantially in magnitude, cell-type specificity and donor consistency, and that clozapine stands out among the pharmacological agents for a response concentrated in sterol and lipid metabolic programs.

### Clozapine regulates astrocytic cholesterol biosynthesis and lipid export pathways

Clozapine’s transcriptional impact was concentrated in steroid biosynthesis in both astrocytes and neurons (Figure 2a). Efavirenz also showed elevated concentration in astrocytes but not in neurons. Azidothymidine showed a similar response in both neurons and astrocytes. When we directly compared the gene-wise responses between the two antipsychotic medications, clozapine elicited substantially stronger induction of *SREBF2*-target cholesterol biosynthesis genes in both neurons and astrocytes compared with haloperidol (Figure 2b). Clozapine and haloperidol have both been associated with dopamine-receptor antagonism ^22^. *DRD2* transcripts were detected in 35 of 37 neuronal donor aggregates and 9 of 17 astrocyte donor aggregates from 24-hour vehicle controls, with median expression of 6.62 and 0.051 counts per million, respectively (Extended Data Figure 2a). The very low astrocytic *DRD2* expression, together with the weaker cholesterol-pathway response to haloperidol, suggests that *DRD2* antagonism alone may not explain clozapine’s effects in our study.

**Figure 2.**
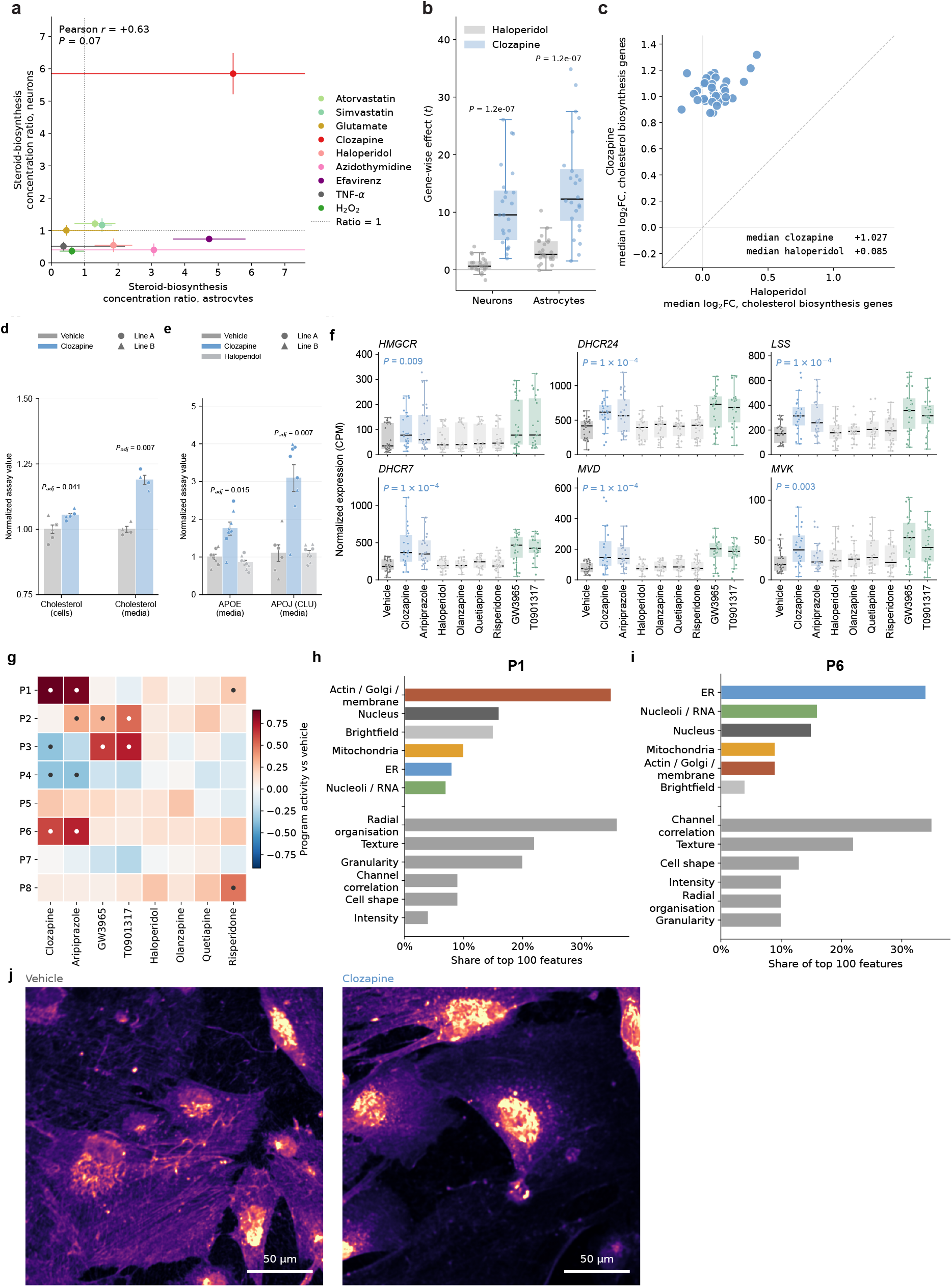
Clozapine engages the astrocyte mevalonate–cholesterol pathway and reorganises astrocyte organelle morphology. **(a)** Steroid-biosynthesis concentration ratios in astrocytes and neurons. Points show treatment estimates with 95% confidence intervals; dotted lines mark 1. Pearson’s *r* and *P* value are shown. **(b)** Gene-wise effects (*t* statistics) on cholesterol-biosynthesis genes under clozapine and haloperidol. Points represent genes; boxes show medians and interquartile ranges. *P* values are from paired two-sided Wilcoxon signed-rank tests. **(c)** Per-donor median log_2_ fold-change across cholesterol-biosynthesis genes under haloperidol and clozapine. Points represent donors; dashed line denotes equal responses. Treatment medians are shown. **(d)** Total cholesterol in astrocytes and conditioned medium after 72 h of clozapine treatment (800 ng/mL). Values were background-corrected and normalized to the pooled control mean. Bars show means *±* SEM; circles and triangles identify wells from two donor lines. Each line contributed one differentiation, with six wells per treatment for intracellular cholesterol and four for conditioned medium, divided equally between lines. **(e)** APOE and APOJ (CLU) in conditioned medium after 72 h of treatment, measured by ELISA. Normalization and symbols follow (d). APOE: eight wells per treatment; APOJ: six vehicle wells and eight per drug treatment, equally divided between lines. In (d,e), displayed *P* values are donor-line-adjusted, two-sided tests with Holm correction. **(f)** Expression of six mevalonate–cholesterol genes in astrocytes exposed to vehicle, six antipsychotics or the LXR-agonists GW3965 and T0901317. Points represent RNA-seq libraries; boxes show medians and interquartile ranges. Wilcoxon rank-sum *P* values compare clozapine with vehicle, adjusted across six genes. **(g)** Eight NeuroPainting morphology programs identified by non-negative matrix factorisation. Colors show vehicle-relative activity adjusted for cell density and exposure duration; dots indicate FDR < 0.05. **(h,i)** Composition of programs P1 (h) and P6 (i), showing percentages of their 100 highest-loading features by imaging channel and measurement family. **(j)** Representative actin/Golgi/membrane-channel images of vehicle- and clozapine-treated astrocytes at 48 h, stained with phalloidin and WGA. Images use identical display ranges. Scale bars, 50 µm.

Clozapine induced systematic upregulation of cholesterol biosynthesis genes across individual cell lines from 34 donors (Figure 2c). In qPCR measurements from two cell lines per comparison, *HMGCR* expression increased in both human primary and iPSC-derived astrocyte lines after 24 h at 800 ng/mL clozapine (unadjusted P = 0.040 and 0.055, respectively; Extended Data Figure 2b,c). *HMGCR* also increased in astrocytes from two iPSC lines after 90 min at 400 ng/mL (P = 0.077; Extended Data Figure 2d). At 24 h, *MVD* and *SCD* expression increased in both lines (P = 0.035 and 0.135, respectively; Extended Data Figure 2e,f).

Additionally, clozapine increased intracellular and extracellular cholesterol (measured by fluorescence), together with APOE and CLU in conditioned medium (measured by ELISA), in astrocytes from both lines (Figure 2d,e). Relative to each line’s vehicle mean, intracellular cholesterol increased by 3.7-7.2%, extracellular cholesterol by 15.5-22.4%, APOE by 1.71-1.80-fold and CLU by 2.51-3.08-fold. Line-adjusted comparisons supported increases in all four assays (Holm-adjusted P < 0.041; Methods).

We next extended these perturbations to four more commonly prescribed antipsychotics, quetiapine, aripiprazole, olanzapine and risperidone, and two LXR agonists, GW3965 and T0901317, with established roles in cholesterol efflux ^23–25^. Antipsychotics were tested across a range of nominal culture concentrations and informed by clinical dosing literature^26,27^ (Methods). A third LXR agonist, LXR-623, was cytotoxic at the concentrations tested and was excluded from analysis (Methods). We measured transcriptional and morphological responses in iPSC-derived astrocytes using bulk RNA-seq and NeuroPainting, an image-based profiling assay we previously developed ^28^.

Clozapine increased expression of all six mevalonate–cholesterol pathway genes examined (Figure 2f). Expression increased 2.45-fold for *MVD*, 2.23-fold for *DHCR7*, 1.84-fold for *LSS*, 1.75-fold for *MVK*, 1.61-fold for *HMGCR* and 1.60-fold for *DHCR24* relative to vehicle. The LXR agonists GW3965 and T0901317 increased expression of the same six genes by an average of 2.06-fold and 1.96-fold, respectively (Figure 2f). Among the other antipsychotics, aripiprazole increased five of the six genes at nominal significance (*HMGCR*, P = 0.014; *DHCR24*, P = 0.002; *LSS*, P = 0.006; *DHCR7* and *MVD*, P < 0.001). Haloperidol, olanzapine and risperidone did not significantly alter expression of these genes, whereas quetiapine reached nominal significance for *MVK* alone (P = 0.047). Among the antipsychotics, coordinated induction of these pathway genes was therefore concentrated in clozapine- and aripiprazole-treated astrocytes at the concentrations and durations we tested.

We next examined the morphological responses to these compounds. NeuroPainting combines six fluorescent stains to visualize nuclei, nucleoli/cytoplasmic RNA, endoplasmic reticulum, actin, Golgi/plasma membrane and mitochondria^28^. Automated multichannel fluorescence imaging and CellProfiler analysis generate quantitative profiles of cell and organelle morphology, including size, shape, staining intensity, texture, radial organization and correlations between channels. Non-negative matrix factorization of NeuroPainting profiles identified eight morphology programs (Methods; Figure 2g; Supplementary Table 5). Clozapine and aripiprazole increased both P1 and P6. P1 was characterized predominantly by actin/Golgi/membrane (AGP) channel features and radial organization measurements (Figure 2h), whereas P6 was characterized predominantly by ER channel features and correlations between organelles (Figure 2i). Risperidone produced a smaller significant increase in P1, but no other compound significantly increased P6. Neither LXR agonist significantly altered P1 or P6, despite inducing the same cholesterol-pathway genes. Thus, clozapine and aripiprazole shared a morphological response distinct from that of the LXR agonists. Representative images illustrate the AGP staining patterns in vehicle- and clozapine-treated astrocytes (Figure 2j).

Together, these findings show that clozapine induces cholesterol-pathway transcription alongside increased cholesterol abundance and altered astrocyte morphology. We next asked whether this response overlaps with the astrocytic cholesterol program induced by neuronal contact.

### Clozapine shares astrocyte responses to neuronal co-culture, with strong induction of cholesterol-related genes

When cultured with neurons, mouse astrocytes upregulate cholesterol biosynthesis and synaptic cell adhesion-associated genes, including *NRXN1* and *NLGN1*, linking astrocyte lipid metabolism with neuron–astrocyte interactions^1,2^. Clozapine’s induction of cholesterol biosynthesis genes in our iPSC-derived astrocytes prompted us to ask whether it also induces astrocytic genes that respond to co-culture with neurons. We compared how iPSC-derived astrocytes respond to clozapine with how mouse astrocytes respond to co-culture with neurons utilizing data from two studies we previously described: the mouse glial/human iPSC-derived neural co-culture versus iPSC-derived neural monoculture contrast reported by Pietiläinen et al. and time-course resolved mouse glial/human iPSC-derived neural co-cultures from Vartiäinen et al. We matched genes through one-to-one mouse–human orthologues, and genes induced in astrocytes cultured with neurons were defined independently of the drug response as those with positive fold-change and FDR < 0.05 (Methods). In the Vartiäinen time course response, 58.8% of the 2,103 co-culture-induced genes had positive fold-changes under clozapine, compared with 42.7% of the remaining 7,075 genes. The Pietiläinen co-culture vs mono-culture showed the same pattern: 54.9% of 2,205 induced genes increased under clozapine, compared with 44.6% of the remaining 7,329 genes (Figure 3a). The corresponding directional odds ratios were 1.92 and 1.51 (Methods; Figure 3 Source Data). Restricting both comparisons to the common gene space produced similar results. Transcriptional similarity with clozapine was stronger the longer astrocytes and neurons were cultured together (Extended Data Figure 2g).

**Figure 3.**
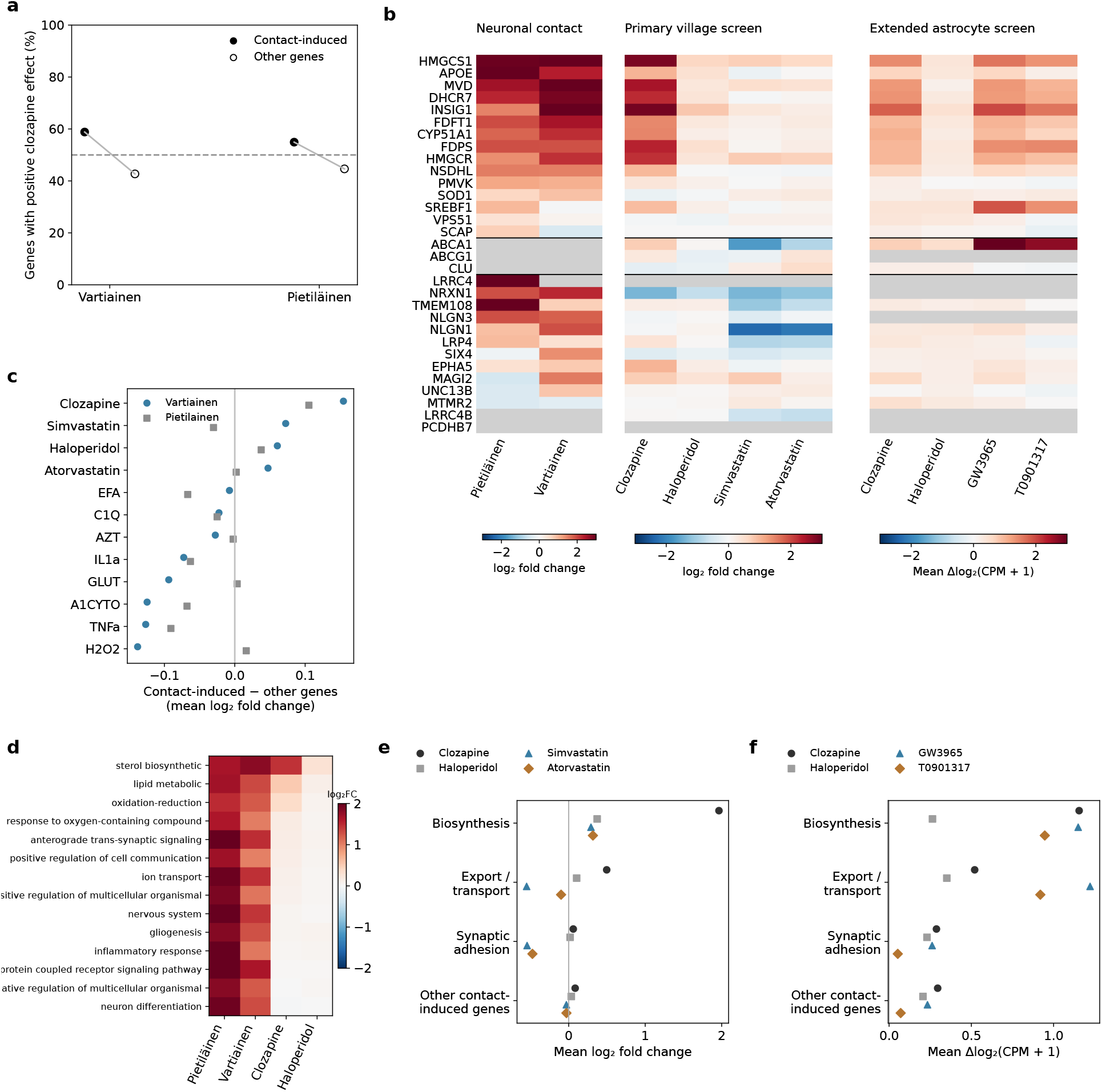
Cholesterol-active compounds share components of the astrocyte response to neuronal contact. **(a)** Percentage of genes with positive clozapine log_2_ fold-change among co-culture-induced genes (filled circles) and remaining genes (open circles), using the Vartiainen and Pietiläinen signatures. The dashed line marks 50%. **(b)** Gene-level responses for published cholesterol-related and synaptic-associated sets, together with *ABCA1*, *ABCG1* and *CLU*. Columns show neuronal-contact comparisons, the primary village screen and the separate extended astrocyte screen. Contact and primary-screen values are log_2_ fold changes; extended-screen values are mean changes in log_2_(CPM + 1) relative to time-matched vehicles. Grey indicates unavailable measurements; colors saturate at *±*3. **(c)** Mean log_2_ fold-change among co-culture-induced genes minus that among remaining genes across twelve perturbations, using the Vartiainen (blue circles) and Pietiläinen (grey squares) signatures. **(d)** Mean log_2_ fold-change across fourteen functional categories in the two contact comparisons and under clozapine and haloperidol. **(e,f)** Mean responses within biosynthesis, transport/export, synaptic-associated and other contact-induced gene sets in the primary village **(e)** and extended astrocyte **(f)** experiments. Points show gene-set means on the respective scales used in (b). Gene sets and measured membership are specified in Methods and Figure 3 source data.

The shared response in astrocytes between clozapine treatment and neuronal co-culture included a strong upregulation of cholesterol-related genes. Within the 15-gene cholesterol set from Pietiläinen et al, 11 genes exceeded +0.5 log_2_ fold-change under clozapine, compared with two under haloperidol (Figure 3b). *HMGCS1*, *INSIG1* and *HMGCR* increased by +2.83, +2.88 and +2.20 log_2_FC units under clozapine, respectively, compared with +3.46, +3.11 and +2.18 in the Vartiäinen co-culture data. The cholesterol-related set also showed larger positive co-culture differential-expression statistics at later durations (Extended Data Figure 2h). The synaptic cell adhesion-associated gene changes from the neuronal contact response were weaker. *NLGN1* increased by +0.89 and +1.94 log_2_ units in the Pietiläinen and Vartiäinen signatures, respectively, but had an estimate near zero under clozapine (*−*0.02). *NLGN3* likewise increased in both co-culture signatures (+1.94 and +1.79) but showed little change under clozapine (*−*0.08). *NRXN1* had a negative clozapine estimate (*−*1.37).

When we examined the neuronal contact response beyond the cholesterol pathway genes, the change following clozapine treatment was modest. Among non-cholesterol related genes induced in the Vartiäinen signature, the median log_2_ fold-change was +0.046. To compare perturbations, we calculated the mean log_2_ fold-change of each co-culture-induced set relative to the remaining genes under each treatment. Clozapine had the largest positive point estimate in both signatures (+0.154 for Vartiäinen and +0.105 for Pietiläinen), whereas the ordering of the other perturbations differed between signatures (Figure 3c). Haloperidol also produced positive shifts in both comparisons with reduced magnitude, while several other treatments differed in direction between signatures.

When we compared the different functional categories reported to be impacted by co-culture with neurons, the largest magnitude of changes were in cholesterol-related processes. Among fourteen GO categories derived from the Pietiläinen et al’s enrichment analysis, the mean clozapine log_2_ fold-change was +1.44 for sterol biosynthesis, +0.52 for lipid metabolism and +0.37 for oxidation– reduction, compared with at most +0.16 for the remaining categories, including those associated with synaptic and neuronal functions (Figure 3d). Directional overlap nevertheless extended beyond cholesterol-related genes: excluding either the published cholesterol set or a broader sterol gene set from the entire analysis universe changed the overlap rates little (Methods). Thus, clozapine shared a modest directional response across a broad set of co-culture-induced genes, with larger induction among cholesterol-related genes and incomplete reproduction of the synaptic-associated response.

We next compared these responses with other cholesterol-active compounds. In the primary village screen, clozapine, simvastatin and atorvastatin all increased biosynthesis-gene expression, with a larger mean increase under clozapine, whereas transport/export and synaptic-associated responses differed among compounds (Figure 3b,e). In the separate extended astrocyte screen, GW3965 and T0901317 produced biosynthesis responses comparable to clozapine and larger mean transport/export responses (Figure 3b,f). Thus, induction of the cholesterol-related component of the contact response was shared across compounds, while the broader expression patterns varied.

### Clozapine engages coordinated neural programs and alters neuronal network activity

We next asked whether clozapine induces gene expression responses in both cell types and neuronal activity in co-cultures.

We generated co-cultures of mouse astrocytes and human excitatory neurons and exposed them to the same panel of antipsychotic and LXR treatments as before (Figure 4a; Supplementary Table 6). Species-specific reads resolved the independent astrocytic and neuronal responses. A latent factor analysis identified two gene expression programs with higher scores under clozapine than under the other treatments (Figure 4b,c). Gene-set enrichment summaries of the clozapine enriched factors associated LF_9 with astrocytic sterol and lipid biosynthesis, and LF_11 with synaptic vesicle exocytosis and neurotransmitter secretion (Figure 4d,e).

**Figure 4.**
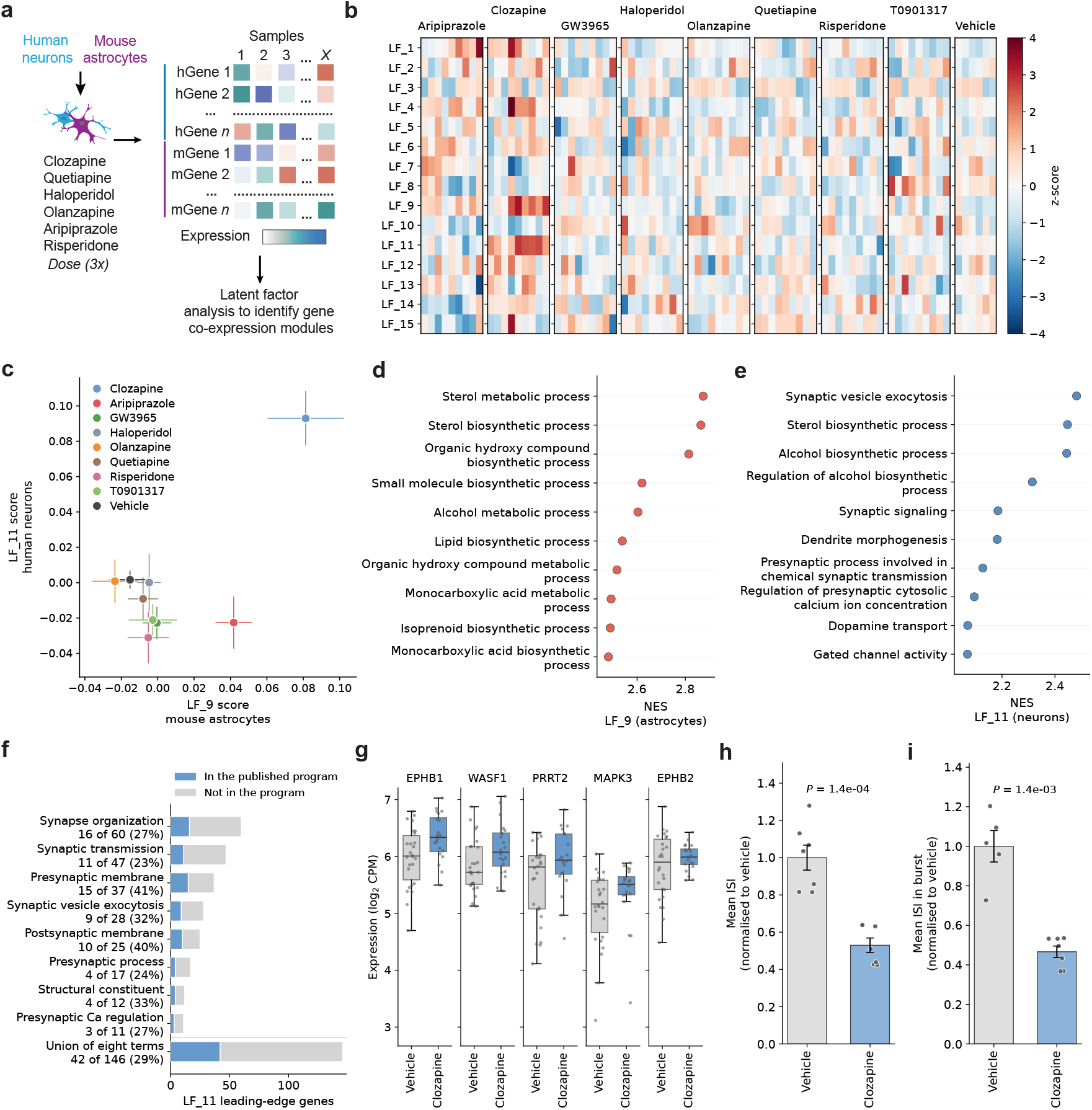
Clozapine engages coordinated neural programs and alters neuronal network activity. **(a)** Schematic of human neuron–mouse astrocyte co-culture and antipsychotic treatment across multiple doses and durations. **(b)** Latent-factor scores across 78 treatment-dose-time conditions. Columns show median scores across library replicates, standardized within each factor. **(c)** Mean LF_9 and LF_11 scores by treatment. Bars show means *±* SEM across conditions. **(d,e)** Ten terms with the highest positive normalized enrichment scores (NES) for the mouse-astrocyte LF_9 (d) and human-neuron LF_11 (e). **(f)** Overlap between leading-edge genes from LF_11-associated synaptic terms and the published neuronal synaptic program associated with astrocyte co-culture. Blue indicates shared genes; grey indicates remaining genes. Labels show shared and total counts and percentages. The bottom row shows the non-redundant union: 42 of 146 genes (28.8%). **(g)** Neuronal expression of five shared genes, *EPHB1*, *WASF1*, *PRRT2*, *MAPK3* and *EPHB2*, under vehicle and clozapine. Points represent samples pooled across doses and durations (26 vehicle; 20 clozapine); boxes show medians and interquartile ranges. Responses across all 42 shared genes are shown in Extended Data Figure 2i. **(h,i)** Multi-electrode array measurements of mean inter-spike interval (ISI; h) and ISI within bursts (i), normalized to vehicle. Bars show means *±* SEM; points represent wells from one cell line and differentiation. Mean ISI: seven wells per treatment; burst ISI: five vehicle and seven clozapine wells. Displayed *P* values are from two-sided Welch tests.

Because astrocytic cholesterol supports synapse formation and function^6,29^, we compared the LF_11 synaptic leading-edge genes with the neuronal contact program. Within the shared SynGO-annotated background, 42 of 146 leading-edge genes belonged to that program (28.8%, Figure 4f; Supplementary Table 7). *EPHB1*, *WASF1*, *PRRT2*, *MAPK3*, and *EPHB2* illustrate shared members in the co-culture expression data (Figure 4g). The complete shared-gene comparison shows heterogeneous drug responses (Extended Data Figure 2i-l).

Next, we asked whether these transcriptional responses were accompanied by altered neuronal activity. In co-cultures exposed to clozapine for 24h, the mean inter-spike interval and interval within bursts were approximately half that of the vehicle controls (Figure 4h,i; 7 vehicle and 7 clozapine wells for mean interval; 5 and 7 for within-burst interval). Both differences were supported by well-level Welch tests after correction across the two measurements (Holm-adjusted P = 2.9 x 10^-4^ and 1.4 x 10^-3^, respectively).

Together, these data show that clozapine engages both an astrocytic sterol and lipid program and a neuronal synaptic program in co-culture, and that this transcriptional response is accompanied by faster neuronal firing.

### Baseline expression is associated with variation in cholesterol-pathway responses following treatment

Having observed the transcriptional and functional effects of clozapine, we next examined the extent to which individuals varied in their cholesterol response. We profiled donor-resolved responses to clozapine, haloperidol and the LXR agonist GW3965 in one expanded donor village (N=107 donors) differentiated into excitatory neurons using scRNA-seq (Figure 5a, Supplementary Table 2). The analyses below contain overlapping donor subsets from this same village, rather than separate villages, as treatment arms with fewer sequencing libraries captured fewer donors (107 donors contributed to baseline-clozapine expression analyses; the treatment-response coefficient matrices contained 86 donors, and the individual-gene between-drug analysis used 88 matched donors). Across the 86 donors, clozapine increased the mean expression of the 25 cholesterol-pathway genes in 85 donors. Median per-gene directional agreement was 94.2%, compared with 59.3% genome-wide (Figure 5b). Mean pathway responses ranged from *−*0.029 to 1.754 log_2_ fold change relative to untreated baseline (Figure 5c). Matched comparisons showed broader increases across biosynthetic genes under clozapine, prominent GW3965 responses in *ABCA1*, *ABCG1* and *SREBF1*, and smaller overall responses to haloperidol (Extended Data Figure 3a–d).

**Figure 5.**
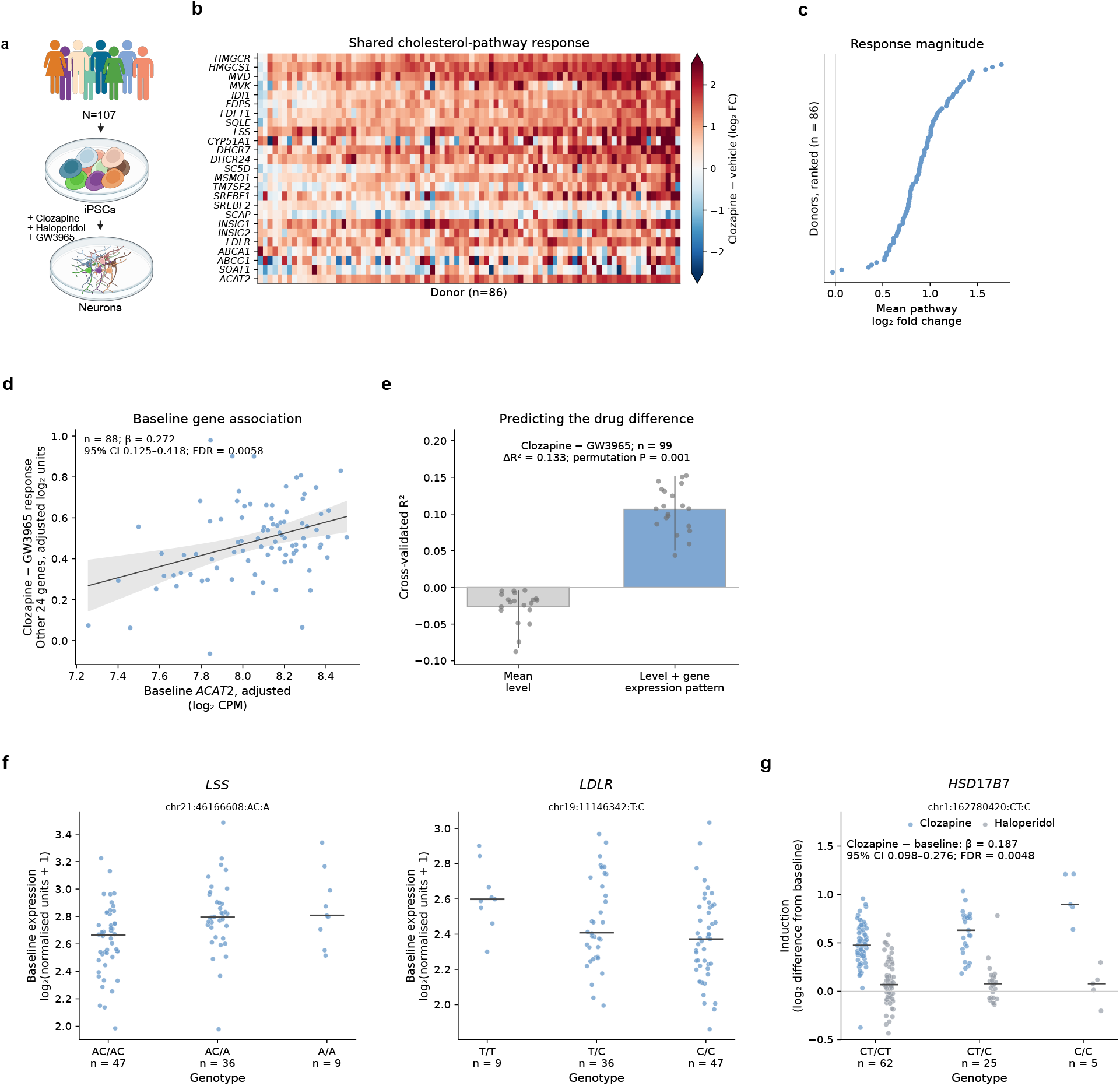
Baseline expression and genetic variation are associated with individual cholesterol-pathway responses. **(a)** Expanded neuronal screen. Wells from one pooled village were treated for 72 h with clozapine (800 ng/mL), haloperidol (1 µM) or GW3965 (50 ng/mL), or left untreated. Donor subsets overlap across analyses. **(b)** Clozapine-induced log_2_ fold-changes across 25 cholesterol-pathway genes in 86 donors, ordered by mean pathway response. Colors saturate beyond *±*2.5. **(c)** Mean cholesterol-pathway log_2_ fold-change for each donor, ranked by response. The vertical line marks zero. **(d)** Untreated *ACAT2* expression versus the clozapine–GW3965 expression difference across the remaining 24 pathway genes in 88 donors, adjusted for their mean baseline expression. The line and shading show the fitted relationship and 95% confidence interval (*β* = 0.272; 95% CI, 0.125–0.418; FDR = 0.0058). **(e)** Cross-validated prediction of the clozapine–GW3965 pathway-response difference in 99 donors using mean baseline pathway expression alone (grey) or with individual-gene expression patterns (blue). Bars show mean R^2^ across 20 repeated five-fold partitions; points show individual repeats and whiskers their 2.5th–97.5th percentiles. Annotations report the prediction gain and permutation *P* value. **(f)** Baseline *LSS* and *LDLR* expression by genotype at chr21:46166608:AC>A and chr19:11146342:T>C (GRCh38). Points represent 92 donors; horizontal lines mark medians. Genotype-group sizes are shown below the axes. **(g)** Clozapine- and haloperidol-induced *HSD17B7* expression changes by genotype at chr1:162780420CT>C in 92 donors. Horizontal lines mark medians. The adjusted clozapine–baseline effect was 0.187 log_2_ units per C allele (95% CI, 0.098–0.276; FDR = 0.0048).

We next asked whether baseline expression was associated with differential responses to cholesterol-related perturbations. Higher measured baseline pathway expression was associated with smaller treated-minus-baseline changes (Extended Data Figure 3e-g).

We examined whether individual genes were associated with differential responses to clozapine and GW3965. For each gene, we modelled the expression difference between the clozapine and GW3965 treatments across the remaining 24 pathway genes, adjusting for their average baseline expression. In the 88 matched donors, baseline expression of 11 genes was associated with this difference after correction across the 25 genes tested (Extended Data Figure 4a; Supplementary Table 8). Associations were positive for genes including *MVD*, *HMGCR*, *ACAT2* and *MVK*, and negative for genes including *SREBF2*, *HMGCS1* and *FDFT1*. Eight associations retained FDR < 0.05 with robust standard errors (Supplementary Table 8).

Higher baseline *ACAT2* expression was associated with a greater pathway response under clozapine relative to GW3965 (*β* = 0.272; 95% CI, 0.125–0.418; FDR = 0.0058; Figure 5d). This association remained positive and significant after omission of each donor in turn and was supported by robust standard errors and donor bootstrapping (Extended Data Figure 4c,d). We also compared clozapine and haloperidol, which yielded no gene-level associations at FDR < 0.05 (Extended Data Figure 4a).

We then tested whether the baseline gene-expression pattern improved prediction beyond the pathway’s average expression level. In 99 donors with baseline, clozapine and GW3965 measurements, adding the individual-gene expression pattern increased mean cross-validated R^2^ for the clozapine– GW3965 pathway-response difference from *−*0.026 to 0.106 (ΔR^2^ = 0.133; permutation P = 0.001; Figure 5e). For the clozapine–haloperidol contrast, the gain was smaller and the final R^2^ remained slightly below zero (*−*0.025 to *−*0.002; ΔR^2^ = 0.023; permutation P = 0.029; n = 92; Extended Data Figure 4b).

These findings associate baseline gene-expression patterns with differential cholesterol-pathway responses to clozapine and GW3965, motivating us to ask whether common genetic variation also contributes to baseline expression differences.

### Common genetic variation is associated with cholesterol-gene expression and the HSD17B7 response to clozapine

The associations between untreated gene-expression and drug-response differences prompted us to ask whether common genetic variation in our donors underlies the expression of the cholesterol pathway genes. We analysed cis-eQTLs in human neurons under baseline, clozapine and haloperidol conditions. Comparing the same 13,331 tested genes in 92 donors represented in all three conditions identified 1,043, 505 and 419 cis-eGenes, respectively (Extended Data Figure 5a, Supplementary Table 9). Our baseline expression-QTL associations agreed in direction with GTEx v8^30^ brain eQTLs in 91.9% of 1,610 matched gene–variant–tissue comparisons spanning 556 genes (Extended Data Figure 5b), as well as 99.6% across 1,952 gene–variant–context comparisons spanning 472 genes in an independent iPSC-derived neuronal dataset of Liang et al. ^31^ (Extended Data Figure 5c).

When examining the 25-gene cholesterol-pathway set above, *LSS* and *LDLR* carried baseline cis-eQTLs in the expression-QTL catalogue (FDR = 0.014 and 0.0024, respectively; Figure 5f). Both genes increased in expression in all but one donor following clozapine treatment. Baseline cis-eQTLs were also detected for the cholesterol and lipid-related genes *HSD17B7* and *TMEM97*, which lie outside the 25-gene set (FDR = 0.0026 and 4 *×* 10*^−^*^6^, respectively).

We next tested whether genotype was associated with treatment-induced changes in expression at the baseline lead variants of 70 lipid-related genes. *HSD17B7*, which encodes a reductase involved in cholesterol biosynthesis ^32^, showed a larger genetic effect under clozapine. At chr1:162780420:CT>C, each C allele was associated with an additional 0.187 log_2_ units of clozapine-induced expression (95% CI, 0.098–0.276; FDR = 0.0048; Figure 5g; Extended Data Figure 5d, Supplementary Table 10). The per-allele expression effect was 0.211 at baseline, 0.398 under clozapine and 0.197 under haloperidol. The clozapine–baseline association remained significant with heteroskedasticity-robust standard errors (FDR = 0.014) and remained positive after omission of each donor in turn (Extended Data Figure 5e,f).

The direct clozapine–haloperidol contrast was 0.201 log_2_ units per allele (95% CI, 0.100–0.302; FDR = 0.012 across the same 70 genes; Extended Data Figure 5d). With robust standard errors, its FDR was 0.051 (Extended Data Figure 5e). A comparison with GW3965 gave a positive contrast of 0.197 log_2_ units per allele in 88 matched donors (95% CI, 0.081–0.314), but did not pass correction across the 70 genes (FDR = 0.084; Extended Data Figures 5d and 6).

**Figure 6.**
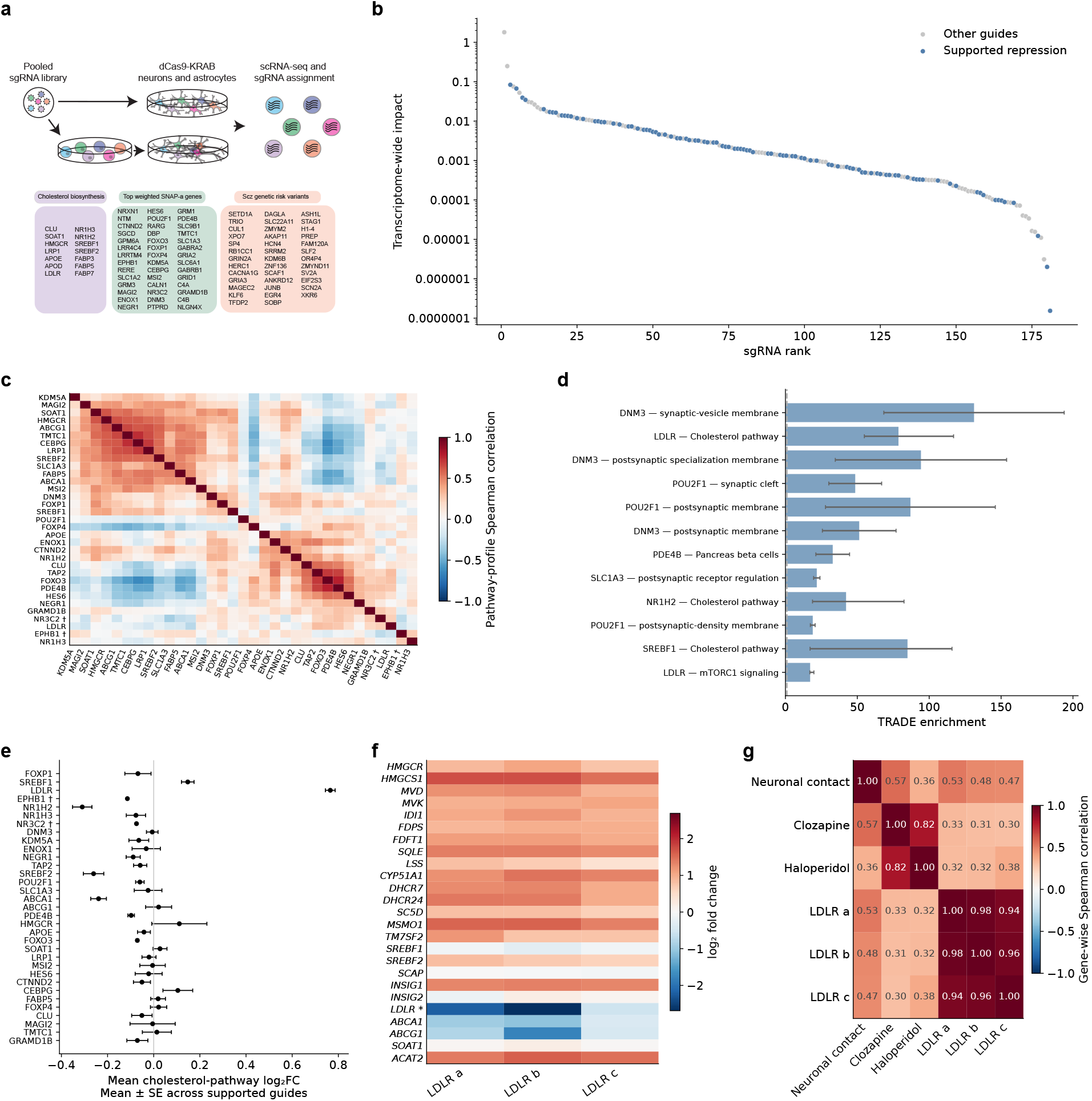
Screen-wide transcriptional impact and the concentrated cholesterol response to *LDLR* repression in astrocytes. **(a)** CRISPRi screen schematic. Pooled neuronal cultures were transduced at the iPSC stage and profiled at D28; pooled astrocytes were transduced at D25 and profiled at D30. **(b)** Ranked transcriptional impact (TI) estimates for 181 astrocyte sgRNA contrasts, colored by evidence for intended-target repression. **(c)** Pairwise Spearman correlations of pathway-response profiles across 877 predefined gene sets among 33 targets with supported repression. Daggers identify targets represented by one supported guide. **(d)** TRADE enrichment for selected target–pathway combinations supported by multiple guides. Enrichment is the fraction of response variance attributed to a gene set divided by its fraction of genes. Bars show medians; whiskers span individual-guide estimates. **(e)** Mean cholesterol-pathway log_2_ fold-change across supported guides for each target. Points show means *±*1 s.e. across guides; daggers indicate single-guide targets. The pathway includes 25 genes, excluding the perturbed gene where applicable. **(f)** Gene-level log_2_ fold-changes across the cholesterol pathway for three supported *LDLR* guides. An asterisk identifies *LDLR*, distinguishing intended-target repression from downstream responses. **(g)** Spearman correlations of signed gene-level response statistics across 22 genes shared among neuronal-contact, clozapine, haloperidol and *LDLR*-repression datasets. Analyses restricted to supported repression include 97 astrocyte guides. Source estimates are provided in Supplementary Table 11.

*TMEM97* expression was associated with genotype under all three conditions. At chr17:28301721:G>A, the per-A-allele expression effects were 0.167 log_2_ units at baseline, 0.323 under clozapine and 0.160 under haloperidol (Extended Data Figure 5g). Its clozapine–baseline response contrast was positive but did not pass correction across the rest of the lipid-gene family (*β* = 0.156; FDR = 0.434; Extended Data Figure 5d).

No association with the aggregate cholesterol pathway-response score survived correction across the 13,066 lead variants tested (Extended Data Figure 5h). Overall, these analyses identify genetic associations with expression at several cholesterol-related genes and clozapine-associated genetic effect at *HSD17B7*.

### CRISPRi reveals diverse transcriptional responses and a concentrated cholesterol response to LDLR repression

Having identified natural genetic associations with cholesterol-gene expression, we next used pooled CRISPRi and single-cell RNA sequencing to test the effects of targeted repression in astrocytes and neurons. The screen included cholesterol-related genes and genes implicated in schizophrenia by genetic or expression studies (Figure 6a). We designed 3-5 sgRNAs per target using CRISPick^33^, including 25 non-targeting control sgRNAs (Methods). SgRNA pools were cloned into the CROP-seq vector ^14^, packaged into lentivirus and delivered to the cells. For the astrocyte experiments, our sgRNA pool was introduced at day 25 and we profiled gene expression five days later. For the neuronal experiments, iPSCs were transduced with the sgRNA pool and then differentiated into excitatory neurons co-culture with mouse astrocytes and profiled at day 28.

We generated pseudobulk gene-expression matrices by summing the individual counts for all cells assigned to a given sgRNA per sequencing library (Methods). Using the pseudobulked data, we computed differential gene expression with DESeq2 ^34^ and assessed intended-target repression for each sgRNA.

Intended-target knockdown was supported for 97 guides targeting 33 astrocyte genes and 133 guides targeting 58 neuronal genes after multiple correction (negative target effect and FDR < 0.05; Extended Data Figures 7a and 9a; Supplementary Tables 11 and 12). We retained all sgRNA contrasts for the screen-wide impact overview and restricted the downstream pathway comparisons to guides with supported repression.

Using the same TRADE framework applied to the pharmacological screen (Figure 1), we found that astrocyte perturbations varied widely in their transcriptome-wide impact (Figure 6b). To examine which targets elicited similar responses, we compared their expression-change profiles across 877 representative gene sets drawn from GO Biological Process and SynGO. These profiles showed shared and opposing patterns amongst the 33 targets with supported repression (Figure 6c; Extended Data Figure 8a,b; Supplementary Table 11).

We explored the downstream agreement between independently designed guides targeting the same gene and its relationship to control-defined expression filtering, transcriptional impact and cell coverage (Extended Data Figure 7b–e). Agreement varied across targets and our focused analyses below examine consistency across guides in the relevant genes and pathways.

We asked which biological pathways our perturbation responses were concentrated in. Enrichment was defined as the fraction of fitted response variance attributable to a set divided by its fraction of measured genes. Among 161 sets examined across all guides, the leading astrocyte target–set pairs included cholesterol-pathway genes and genes annotated to synaptic membranes and the synaptic cleft (Figure 6d; Supplementary Table 11). However, these concentrations were accompanied by different expression patterns. Cholesterol genes increased following *LDLR* and *SREBF1* repression but decreased following *NR1H2* repression, whereas several highly enriched synaptic sets showed modest or mixed mean changes across guides (Extended Data Figure 8c). In the *DNM3* synaptic-vesicle membrane set, different individual genes accounted for approximately two-thirds of the observed squared fold changes in the two guides.

Similar to what we observed with the clozapine response, a substantial fraction of the response to *LDLR* repression was concentrated in cholesterol-pathway genes. The three supported *LDLR* guides had between 194–274 differentially expressed genes passing FDR < 0.05. Although the 24 other cholesterol-pathway genes comprised only 0.19% of the genes we analyzed, they accounted for 10.5–22.4% of the fitted response variance for these guides, a 55–117-fold concentration relative to their representation in the data (Figure 6d; Supplementary Table 11).

We examined the same 25-gene cholesterol-pathway set used above, which includes biosynthetic enzymes, regulatory genes and transporters. Each guide’s own target was excluded from its down-stream pathway score. *LDLR* repression produced the strongest positive mean pathway response among targets (Figure 6e). The strongest negative target averages were observed for *NR1H2* (-0.31 log2FC; five guides), *SREBF2* (-0.26 log2FC; three guides) and *ABCA1* (-0.24 log2FC; three guides) (Figure 6e). Three independently designed guides, a, b and c, reduced *LDLR* expression by 2.22, 2.69 and 0.56 log2FC and increased mean expression of the remaining 24 pathway genes by 0.79, 0.78 and 0.72 log2FC, respectively. Each guide increased expression of 20 of these 24 genes; 18, 17 and 15 increases, respectively, met the original gene-level differential-expression threshold of adjusted P < 0.05 (Figure 6f). All three increased *SREBF2* and *INSIG1*, whereas none significantly altered *SREBF1* after gene-level multiple-testing correction (Figure 6f). Full guide-level cholesterol enrichment estimates are shown in Extended Data Figure 8d. The three supported *LDLR* guides also yielded positive transcriptome-wide TRADE correlations of 0.77–0.96 (Extended Data Fig. 7b; Supplementary Table 11). These estimates provide complementary transcriptome-wide evidence that the response to *LDLR* perturbation was highly consistent across guides.

At single-cell resolution, a cholesterol metagene score, calculated as the mean log1p-normalized expression of the pathway genes excluding *LDLR* (Methods), was higher in cells containing *LDLR*-targeting guides than in non-targeting controls across all sequencing batches from the same pooled culture (Extended Data Figure 7f). Analyses using an expression-matched background adjustment gave consistent results (Extended Data Figure 8e,f; Supplementary Table 11). The background-adjusted increase was present within each of the cellular states defined from control-cell expression for all three guides (Extended Data Figure 8i,j). Across two-, three- and four-state partitions, the contribution from differences in state proportions contributed less than 3% of each guide’s total score difference in absolute magnitude (Extended Data Figure 8k,l).

The *LDLR* response shared induction of cholesterol-pathway genes with the neuronal contact and clozapine responses described above (Figures 3b and 6g). Across the 22 shared downstream genes, Spearman correlations of signed response statistics of the three *LDLR* guide profiles with co-culture were 0.53, 0.48, and 0.47, and with clozapine were 0.33, 0.31, 0.30. Simvastatin and atorvastatin from our initial drug perturbation experiments shared positive expression responses with *LDLR* repression across several biosynthetic genes, including *HMGCR*, *HMGCS1*, *LSS*, *SQLE* and *DHCR24*. Both statins also showed decreased *ABCA1* expression, as observed following *LDLR* repression but opposite to its response under clozapine (Extended Data Figure 8m,n).

Thus, *LDLR* repression induced a coordinated astrocyte cholesterol-pathway response across three supported guides. Its overlap with neuronal contact, clozapine and statins responses was strongest among biosynthetic genes, with distinct responses in cholesterol-export genes such as *ABCA1*.

### Neuronal perturbations reveal shared and opposing pathway responses

We applied the same analytical approach to neurons. The neuronal screen included 133 guides with supported intended-target repression across 58 targets. 43 targets had several supported guides and 15 had one. TRADE transcriptome-wide impact varied across guides and targets (Figure 7a). We retained all supported targets in the screen-wide comparisons and examined guide agreement when selecting examples for closer analysis (Extended Data Figure 9a,b; Supplementary Table 12).

**Figure 7.**
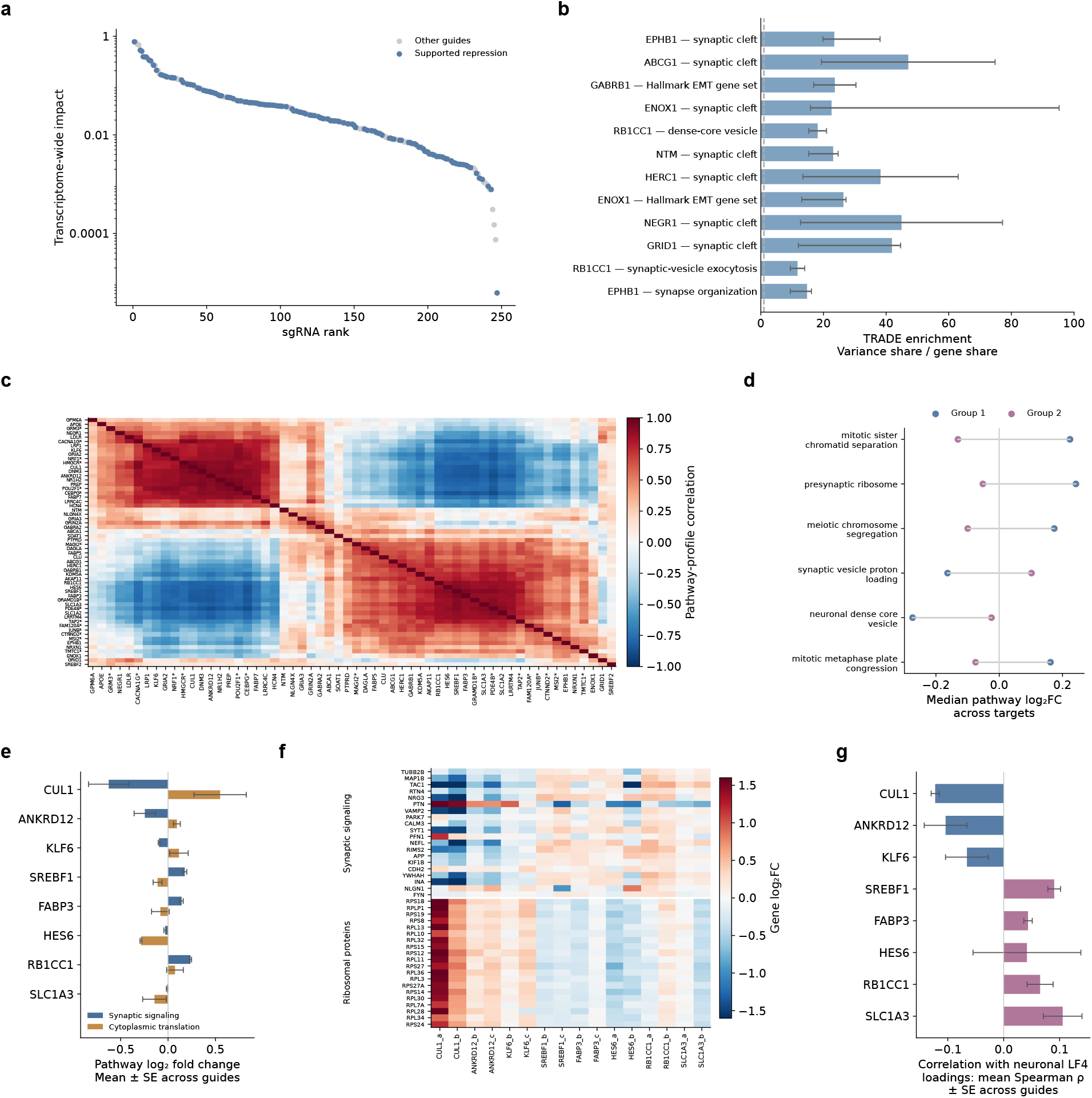
Neuronal perturbations reveal shared and opposing pathway responses. **(a)** Ranked TRADE transcriptome-wide impact across 247 neuronal sgRNA contrasts. Blue identifies 133 guides with supported intended-target repression; grey indicates other guides. The vertical axis is logarithmic. **(b)** Twelve target–pathway combinations with high TRADE concentration across multiple supported guides. Concentration is the fraction of response variance attributed to a gene set divided by its fraction of measured genes. Bars show guide medians; whiskers span individual-guide estimates. **(c)** Spearman correlations of pathway-response profiles across 58 supported targets and 886 representative pathways. Asterisks identify 15 single-guide targets. Clustering identifies two groups summarized in (d). **(d)** Six pathways showing the largest differences in median expression response between the two groups, comprising 27 and 31 targets. Points summarize target-level pathway responses; lines connect group medians. **(e)** Mean log_2_ fold-change in synaptic-signaling and cytoplasmic-translation genes for eight targets whose group membership remained consistent across 200 repetitions of selecting one supported guide per target. Bars show means *±*1 s.e. across two guides per target. **(f)** Gene-level log_2_ fold-changes for the same targets and guides. Each block shows the 20 most highly expressed control genes within the synaptic-signaling or cytoplasmic-translation set. The latter comprises RPL/RPS ribosomal-protein genes. **(g)** Correlations between gene-level log_2_ fold-changes and published neuronal LF4 loadings. Bars show mean Spearman correlations *±*1 s.e. across two guides per target; colors indicate the groups in (d). Individual-guide results are shown in Extended Data Figure 9e,f. Source estimates are provided in Supplementary Table 12.

To identify where responses were concentrated, we examined TRADE estimates for predefined SynGO, Hallmark and cholesterol gene sets. We also included gene sets previously nominated from schizophrenia post-mortem studies^35^ (synaptic-neuron astrocyte program or SNAP genes). Several perturbations showed concentrated effects in synaptic annotations, including synaptic-cleft genes under *EPHB1*, *ABCG1* and *ENOX1* repression, and dense-core-vesicle and synaptic-vesicle-exocytosis genes under *RB1CC1* repression (Figure 7b). Broader pathway responses, concentration estimates and separate GO BP/SynGO companion tests are shown in Extended Data Figure 10a–d (and Supplementary Table 12).

We next compared the relative patterns of expression changes across 886 representative predefined pathways. Target-level profiles showed both shared and opposing responses (Figure 7c). *CUL1* and *ANKRD12* had similar gene-wise responses (Spearman *ρ* = 0.90), whereas *SREBF1*, *FABP3*, *HES6*, *RB1CC1* and *SLC1A3* formed another set of mutually similar profiles (*ρ* = 0.86–0.93). The broad structure was also apparent with alternative pathway summaries (Extended Data Figure 9c). A two-group summary highlighted differences involving synaptic and vesicle-related genes, ribosomal genes and chromosome-related annotations (Figure 7d).

Group assignment varied for many targets when one supported guide per target was sampled and the clustering repeated. *CUL1*, *ANKRD12* and *KLF6* retained one assignment in all 200 guide-choice repeats, while *SREBF1*, *FABP3*, *HES6*, *RB1CC1* and *SLC1A3* retained the other (Extended Data Figure 9d). We examined these targets in greater detail.

Both *CUL1* guides and both *ANKRD12* guides decreased mean synaptic-signaling expression and increased cytoplasmic-translation expression, whereas both *SREBF1* guides showed the opposite pattern (Figure 7e; Extended Data Figure 9e). Other targets varied within these gene programs. For example, *FABP3* repression increased synaptic-signaling expression but decreased translation related genes, whereas *HES6* guides decreased both sets. Gene-level displays showed coordinated increases in the selected ribosomal-protein transcripts under *CUL1* and *ANKRD12* repression and decreases under *SREBF1* and *HES6* repression (Figure 7f).

To relate these responses to the coordinated astrocyte-neuron biology we explored earlier, we compared perturbation responses with the neuronal LF4 loadings underlying the human SNAP program described by Ling et al. ^35^. Correspondence was modest and varied in direction. Both guides targeting each of *CUL1*, *ANKRD12* and *KLF6* had negative correlations, whereas both guides targeting each of *SREBF1*, *FABP3*, *RB1CC1* and *SLC1A3* guides had positive correlations (Figure 7g; Extended Data Figure 9f). Results for all supported neuronal targets are shown in Extended Data Figure 10e. Among 44 eligible genes from the top 50 neuronal LF4 genes, *CUL1* repression decreased expression of 11 and increased four passing the original adjusted P < 0.05 threshold in both guides. Extending the set to the top 100 yielded 84 eligible genes, with 27 decreases and four increases (Extended Data Figure 10g).

Compared with response in neurons exposed to clozapine, we identified opposing changes in ten top-100 LF4 genes. Eight, including *PIK3R1*, *BICDL1* and *BRSK1*, decreased under both *CUL1* guides but increased under clozapine. *RBM3* and *WWC1* showed the opposite pattern (Extended Data Figure 10f). These comparisons passed the original gene-level adjusted P < 0.05 threshold for clozapine and each *CUL1* guide. Together, these observations identify shared and opposing neuronal expression responses to repression of individual cholesterol biosynthesis pathway and schizophrenia-associated genes, including modest overlaps with the human SNAP program. Although *CUL1* repression opposed clozapine at some SNAP genes, no perturbation reproduced or reversed clozapine’s effects or schizophrenia-associated gene expression phenotypes.

## Discussion

By combining pharmacological perturbation across genetically diverse human neural cultures, cis-eQTL mapping under drug exposure and CRISPRi, we identify components of an astrocytic cholesterol-related program that can be engaged pharmacologically and genetically. Clozapine produced a relatively small global transcriptional response but concentrated that response in cholesterol biosynthesis and export. This response corresponded to the cholesterol-related component of the astrocyte response to neuronal contact, with little corresponding change in the astrocytic synaptic cell-adhesion component. Together with increased cholesterol in astrocytes and conditioned medium, these findings identify shared cholesterol-related responses without reproducing the full transcriptional response to neuronal contact.

There is additional evidence that clozapine, a recommended antipsychotic medication for treatment-resistant schizophrenia, is linked to cholesterol regulation ^36^. In human CNS-derived cell lines clozapine and several other antipsychotics activate sterol regulatory element-binding protein (SREBP)-controlled expression of lipid-biosynthesis genes, with marked differences among drugs and cell types ^37^. This transcriptional response has been linked to disruption of intracellular cholesterol trafficking and synthesis^38^, and cholesterol-biosynthesis gene induction has similarly been reported in human induced excitatory neurons exposed to clozapine ^39^. These observations motivated us to examine how clozapine’s cholesterol-related effects relate to astrocyte–neuron expression programs, cholesterol abundance and neuronal activity in human neural cultures.

Clozapine’s strongest correspondence with neuronal contact involved cholesterol-related genes. Haloperidol showed similar genome-wide correspondence with neuronal contact, yet induced substantially smaller changes in cholesterol-pathway expression. Statins also induced biosynthesis genes in the primary screen, while LXR agonists produced biosynthesis responses comparable to clozapine in the extended astrocyte screen. The compounds differed in transport/export responses, including *ABCA1*. These comparisons place clozapine within a broader cholesterol-regulatory response shared by pharmacologically distinct compounds. Among the other antipsychotics, aripiprazole increased a subset of cholesterol genes at the tested exposures. Clozapine’s transcriptional response corresponded more closely to later stages of the neuronal-contact response than to its earliest changes.

Altered intracellular cholesterol trafficking or availability could engage sterol-sensitive SCAP– SREBP regulation ^38,40^, providing one possible explanation for the coordinated transcriptional response, but our experiments do not identify the upstream signal. Clozapine-treated co-cultures also showed shorter inter-spike intervals, providing a functional observation alongside the transcriptional and cholesterol measurements. These recordings do not distinguish direct neuronal effects from effects mediated by astrocytes or establish that cholesterol changes caused the increase in firing rate.

The expanded neuronal village revealed variation in observed response magnitude within a broadly shared cholesterol-pathway response. Baseline expression patterns carried information about differential responses to clozapine and GW3965, and baseline *ACAT2* expression was positively associated with the between-drug response difference across the remaining pathway genes. These observations suggest that baseline transcriptional state is informative about variation in responses to distinct cholesterol-related perturbations.

Common genetic variation contributed to baseline expression differences at several cholesterol-related genes in neurons, including *LSS*, *LDLR*, *TMEM97* and *HSD17B7*, and our cis-eQTL associations showed high directional concordance with external brain expression-QTL datasets, supporting their relevance beyond this experimental system. At *HSD17B7*, the genetic effect on expression increased under clozapine, providing a focused example of clozapine-enhanced genetic regulation. We found no significant associations in the aggregate cholesterol pathway-response, which we were largely underpowered to detect. The results therefore identify genetic contributions to expression at individual pathway-related genes and a specific association with drug-induced expression. Although the baseline-expression and eQTL analyses motivate a connection between genetic background, cellular state and drug response, they do not establish a causal chain linking these observations.

The CRISPRi screen placed these cholesterol-related responses within a broader landscape of perturbation-associated transcriptional changes. Perturbations differed in both their overall transcriptomic impact and the extent to which that impact was concentrated in particular pathways. Within this screen, *LDLR* repression produced a consistent cholesterol-pathway response across supported guides, linking a regulator of cholesterol uptake to the coordinated transcriptional response examined throughout this study.

Three guides repressing *LDLR* induced coordinated increases in cholesterol-biosynthesis genes, together with *SREBF2* and *INSIG1*, whereas repressing *SREBF1* did not. This pattern is consistent with a compensatory transcriptional response to reduced lipoprotein-derived cholesterol uptake ^41,42^. Shared induction of biosynthetic genes after *LDLR* repression, statin treatment and clozapine exposure suggests convergence on cholesterol-regulatory transcription ^43^. They varied in their *ABCA1* responses, however, indicating that these perturbations do not produce an identical lipid-handling program. Thus, *LDLR* provides a genetic entry point into a transcriptional program overlapping that upregulated by clozapine.

The neuronal screen revealed expression responses involving synaptic and translation-related genes. *CUL1* and *ANKRD12* repression reduced synaptic-signaling expression while increasing translation-related expression, and *SREBF1* repression showed the opposite pattern. The broader pathway comparisons revealed similarities across additional targets, with variation in the individual genes and pathways affected. These responses affected genes with high loadings in the published neuronal LF4 expression program, and some of these genes changed in opposite directions following *CUL1* repression and clozapine exposure, identifying specific points of correspondence between genetic and pharmacological perturbations. The modest overall correspondence indicates that this opposition involves a subset of genes rather than reversal of the full program. Together, these analyses connect individual perturbations to components of coordinated neuronal expression and identify candidate responses for further functional investigation. Independent CRISPR-based work in human neurons has likewise linked perturbation of schizophrenia-associated genes to synaptic phenotypes, demonstrating simvastatin exposure partially ameliorated increased synaptic puncta following *TMEM219* activation ^44^. This provides a separate example of an interaction between a genetic perturbation and cholesterol-related pharmacology.

Several limitations define the scope of these conclusions. Nominal drug concentrations in culture cannot be directly equated with clinical exposure, unbound concentrations were not measured, and differences in protein binding limit comparisons with plasma concentrations. The two-dimensional cultures lack microglia, oligodendrocytes and cortical architecture. Cross-experiment correspondence identifies shared downstream transcriptional responses rather than shared upstream mechanisms, and increased expression of biosynthetic genes does not itself establish increased synthesis flux. Whether engagement of astrocytic cholesterol supply contributes to clozapine’s therapeutic efficacy remains to be tested. We show that cholesterol-related transcription can be engaged through pharmacological and genetic perturbations, that observed response magnitudes vary across donors, and that baseline expression and common variation contribute to distinct aspects of this biology. These findings provide a basis for testing how treatment, cellular state and genetic background influence astrocyte–neuron function.

## Extended Data

**Extended Data Figure 1.**
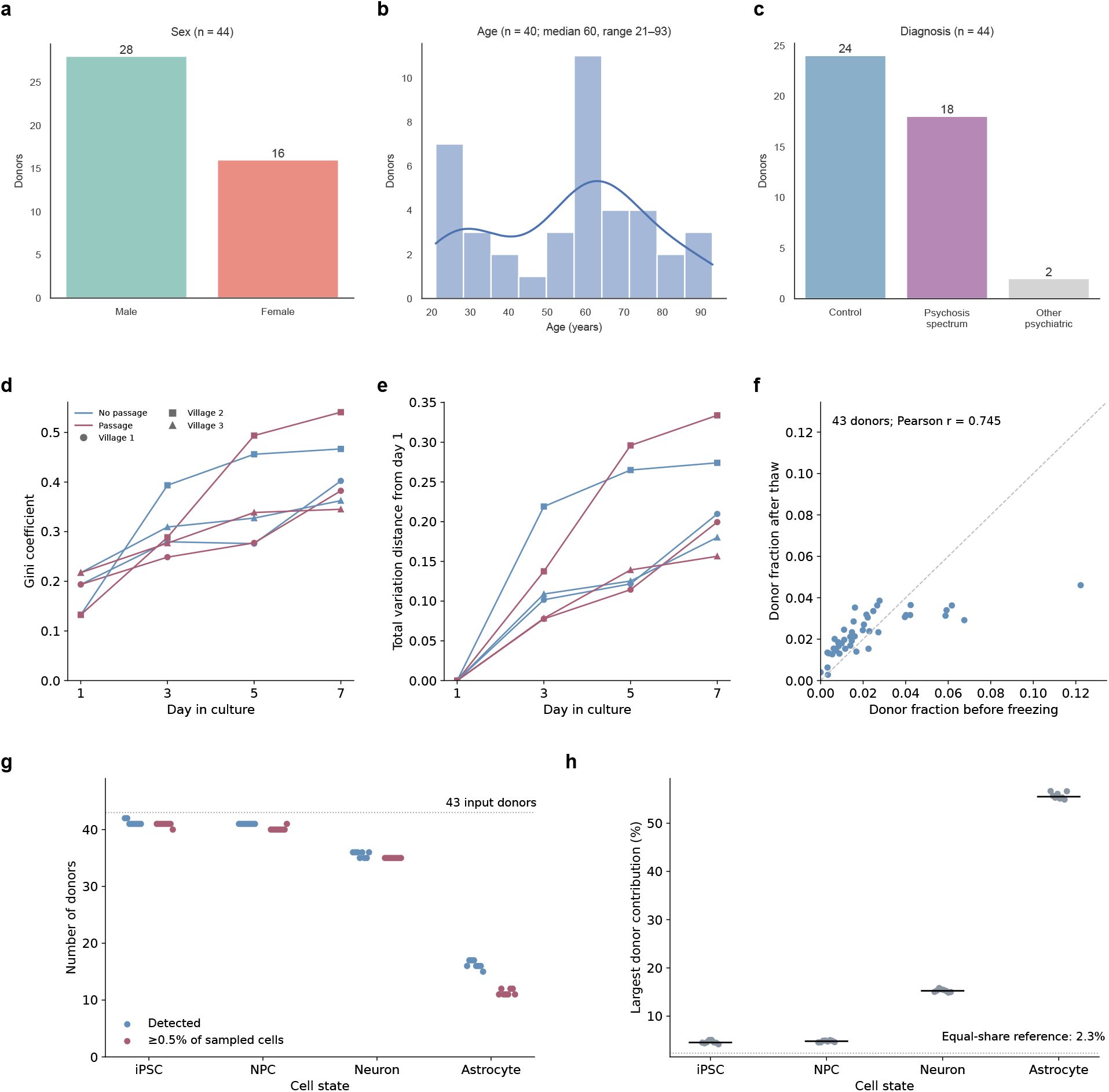
Cohort demographics and donor representation during culture, cryopreservation and neural differentiation. **(a–c)** Recorded sex (a), age (b) and clinical diagnosis **(c)** of the 44-donor primary pharmacological-screen cohort. Age was available for 40 donors (median 60 years; range 21–93). Diagnosis groups comprise 24 controls, 18 psychosis-spectrum donors and two donors with other psychiatric diagnoses. **(d,e)** Gini coefficient of donor representation (d) and total variation distance from day-1 composition (e), measured by Census-seq in three villages containing 45, 46 and 48 donors. Colors indicate handling regimes; symbols indicate villages. (f) Donor fractions before freezing and after thawing in a 43-donor village (Pearson *r* = 0.745). Points represent donors; the dashed line denotes equal fractions. **(g,h)** Donor representation across iPSCs, neural progenitor cells (NPCs), neurons and astrocytes, with eight samples per cell state. Panel g shows the number of detected donors and those contributing at least 0.5% of sampled cells; the dotted line marks the 43 input donors. Panel h shows the largest donor contribution per sample, with horizontal bars indicating medians and a reference marking equal representation (2.3%). Source summaries are provided in Supplementary Table S1.

**Extended Data Figure 2.**
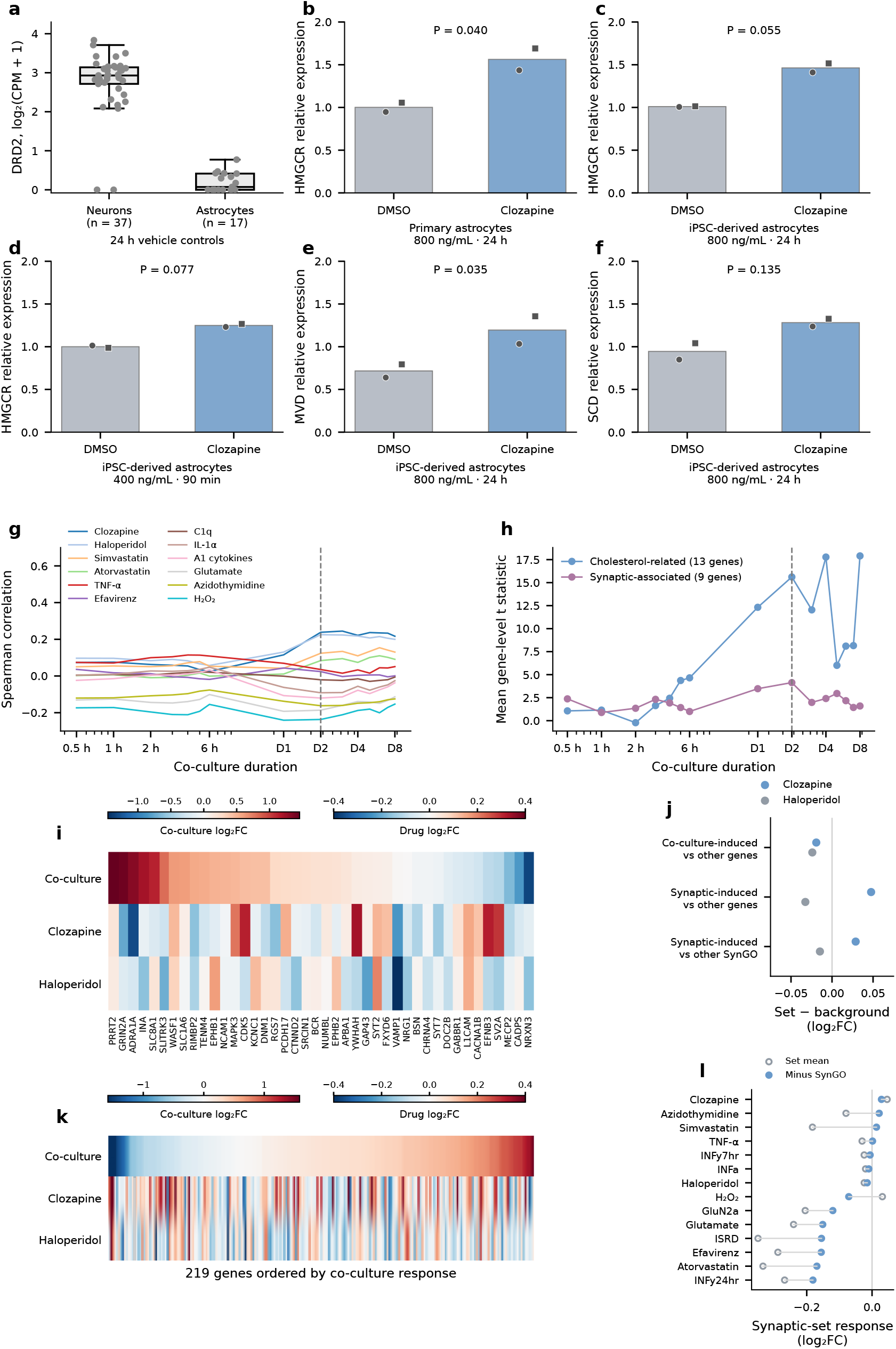
DRD2 expression, qPCR validation and correspondence with neuronal contact programs. **(a)** Donor-level *DRD2* expression in 24 h DMSO controls from neuronal and astrocyte villages, shown as log_2_(CPM + 1). Points represent 37 neuronal and 17 astrocyte donors; boxes show medians and interquartile ranges. **(b,c)** *HMGCR* expression in primary human (b) and iPSC-derived (c) astrocytes after 24 h of clozapine treatment at 800 ng/mL. **(d)** *HMGCR* expression in iPSC-derived astrocytes after 90 min at 400 ng/mL clozapine. **(e,f)** *MVD* (e) and *SCD* (f) expression in iPSC-derived astrocytes after 24 h at 800 ng/mL clozapine. In (b–f), bars show means across two cell lines; symbols show replicate averages for each line. Relative expression was normalized to *GAPDH* and *RPL13* and pooled DMSO controls. Displayed *P* values are from two-sided paired *t*-tests of log_2_-transformed line means. **(g)** Gene-level Spearman correlations between astrocyte drug-response profiles and the published astrocyte co-culture response across 15 co-culture durations. **(h)** Mean differential-expression *t* statistics across 13 cholesterol-related and nine synaptic-associated genes. In (g,h), time axes are logarithmic; dashed lines mark 48 h. **(i)** Co-culture and drug responses of 42 neuronal LF_11 leading-edge genes shared with the published synaptic program. **(j)** Mean drug log_2_ fold-change among co-culture-induced neuronal genes relative to the indicated background, for broad induced and synaptic-induced sets containing 7,025 and 219 genes, respectively. **(k)** Responses of all 219 synaptic co-culture-induced genes, ordered by co-culture response. Heatmaps in (i,k) use separate co-culture and drug color scales. **(l)** Mean synaptic-set response across neuronal perturbations, before and after subtraction of the remaining SynGO-gene mean. Panels j–l use the neuronal village screen. Gene-level results are provided in Supplementary Table 7 and source data.

**Extended Data Figure 3.**
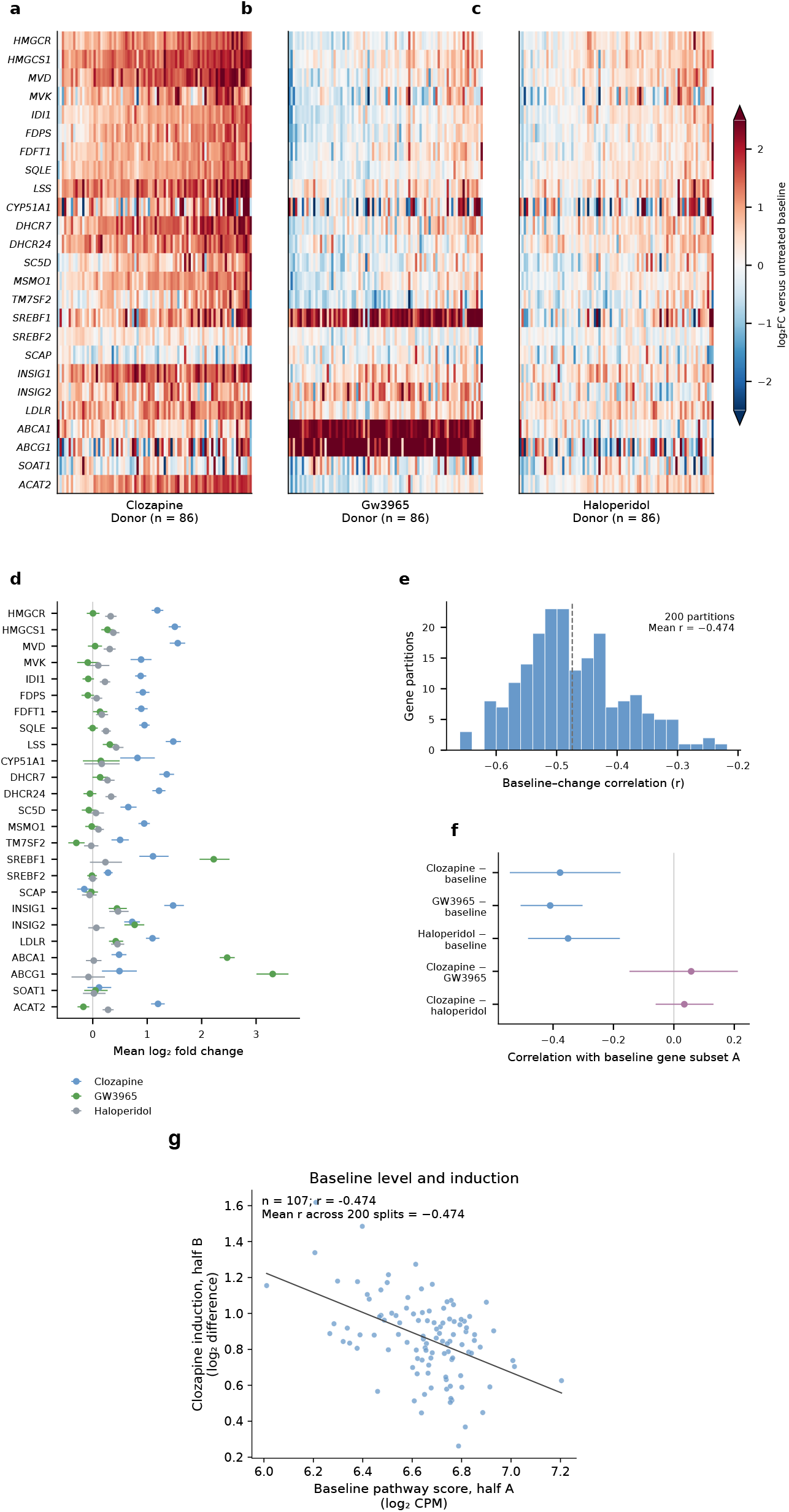
Cholesterol-pathway responses across donors and compounds. **(a–c)** Responses to clozapine **(a)**, GW3965 **(b)** and haloperidol **(c)** across 25 pathway genes and 86 donors. Colors show log_2_ fold-change relative to untreated baseline, saturated beyond *±*2.5. Gene and donor ordering matches Figure 5b. **(d)** Mean gene responses with 95% donor-bootstrap intervals. **(e)** Pearson correlations between baseline expression and clozapine-induced change calculated from complementary pathway-gene subsets in 107 donors, across 200 gene partitions. The dashed line marks the mean correlation (*−*0.474); Panel g shows a representative partition. **(f)** Corresponding analyses in 88 donors measured under all four conditions. Blue indicates treated-minus-baseline responses; purple indicates between-drug differences. Points show mean correlations; bars span the 2.5th–97.5th percentiles across 200 gene partitions. **(g)** Baseline expression versus clozapine-induced change calculated from complementary pathway-gene subsets in 107 donors. Points represent donors; the line shows a linear fit. The displayed partition has the correlation closest to the mean across 200 partitions. Partitioning genes does not remove shared baseline-measurement variation.

**Extended Data Figure 4.**
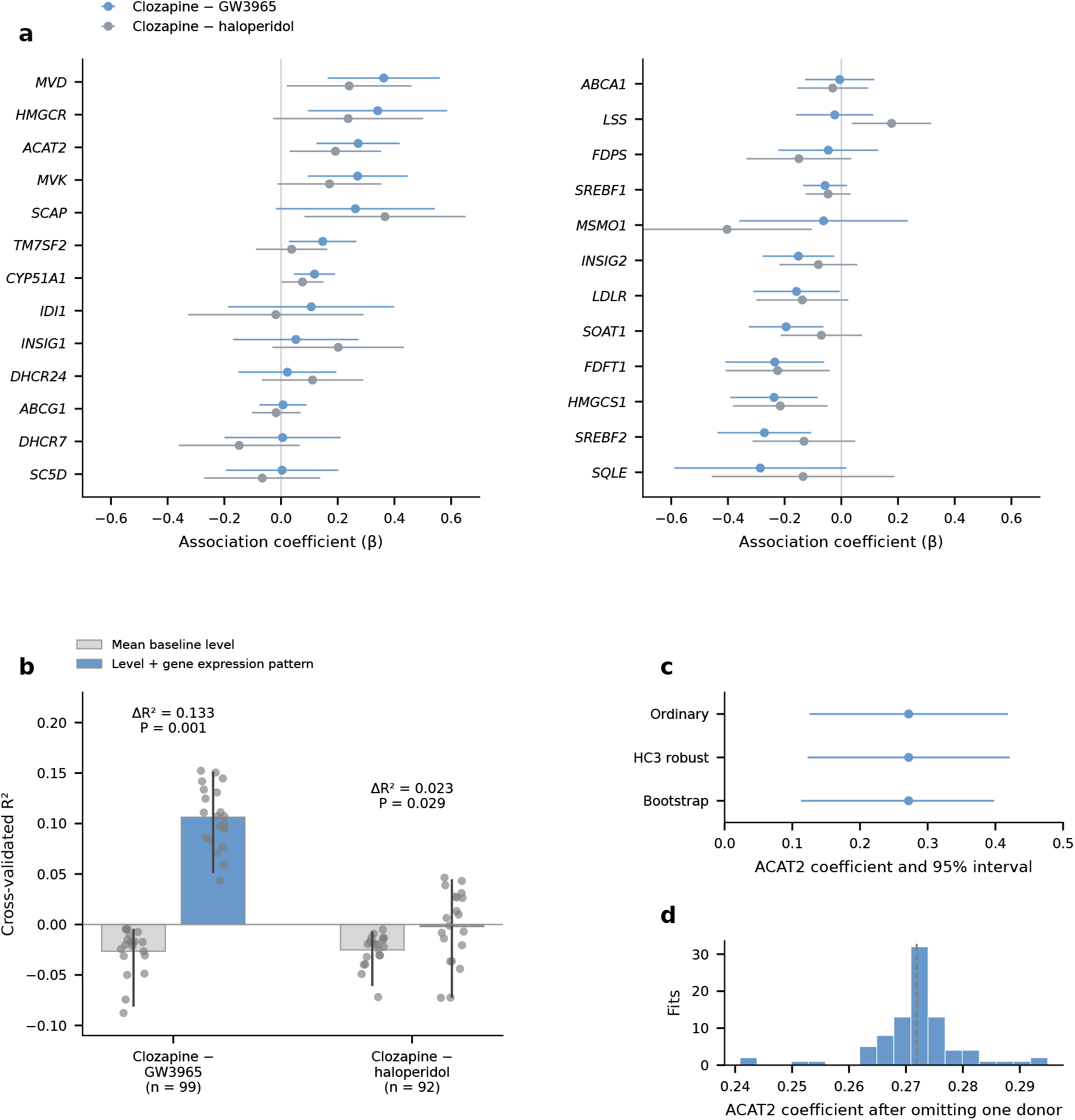
Baseline-expression associations and prediction sensitivity. **(a)** Associations between each gene’s baseline expression and the clozapine–GW3965 (blue) or clozapine–haloperidol (grey) response difference across the remaining 24 pathway genes in 88 donors, adjusted for their mean baseline expression. Points show coefficients; bars indicate 95% confidence intervals. **(b)** Cross-validated prediction of between-drug pathway-response differences using mean baseline expression alone or with individual-gene expression patterns. Bars show mean R^2^ across 20 repeated five-fold partitions; points show repeats and whiskers at their 2.5th–97.5th percentiles. Annotations report prediction gains and permutation *P* values. The GW3965 and haloperidol comparisons include 99 and 92 donors, respectively. **(c)** *ACAT2* association coefficient (*β* = 0.272) with ordinary, HC3 robust and donor-bootstrap 95% confidence intervals. **(d)** *ACAT2* coefficients after omitting each donor individually. The dashed line marks the full-sample estimate. All 88 omission estimates remained positive and passed FDR < 0.05 under ordinary and HC3 inference. Full results are provided in Supplementary Table 8.

**Extended Data Figure 5.**
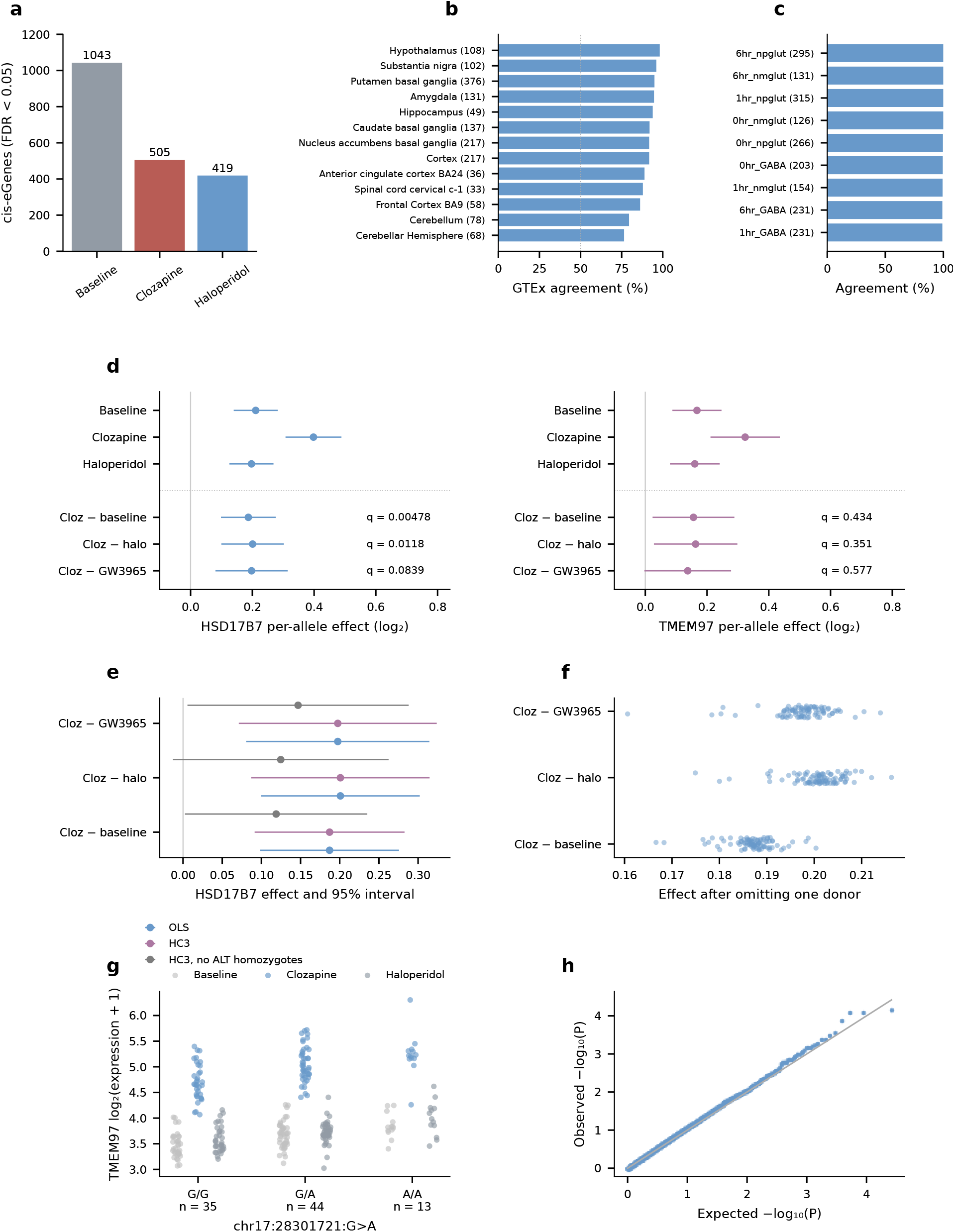
cis-eQTL detection, external agreement and response-effect sensitivity. **(a)** Cis-eGene counts at FDR < 0.05 across 13,331 genes in 92 shared donors: 1,043 at baseline, 505 under clozapine and 419 under haloperidol. **(b)** Directional agreement of baseline associations with GTEx v8 brain eQTLs across 1,610 matched comparisons spanning 556 genes and 13 tissues. Overall agreement was 91.9% (95% interval, 89.2–94.3%). **(c)** Directional agreement with Liang et al. across nine neuronal contexts: 99.6% across 1,952 comparisons representing 472 unique gene–variant pairs (95% interval, 99.2–100.0%). Parentheses indicate comparison counts. Intervals in (b,c) are from gene-clustered bootstrap resampling. **(d)** Per-allele effects at *HSD17B7* chr1:162780420>C and *TMEM97* chr17:28301721>A. Points show estimates with 95% confidence intervals. Within-condition effects appear above the dotted line; clozapine-minus-reference effects appear below. Baseline and haloperidol comparisons include 92 donors; GW3965 includes 88. Annotated *q* values are corrected across 70 lipid-related genes per contrast. **(e)** *HSD17B7* contrast estimates with conventional or HC3 robust standard errors, including analyses excluding alternate homozygotes. **(f)** *HSD17B7* coefficients after omitting each donor individually. All estimates remained positive. **(g)** *TMEM97* expression by genotype in 92 donors, shown as log_2_(normalized expression + 1). Groups comprise 35 G/G, 44 G/A and 13 A/A donors. **(h)** Quantile–quantile plot for associations between 13,066 lead variants and the mean clozapine-induced cholesterol-pathway response in 92 donors. No association passed FDR < 0.05. The diagonal indicates the uniform-*P* reference. Source results are provided in Supplementary Tables 9 and 10.

**Extended Data Figure 6.**
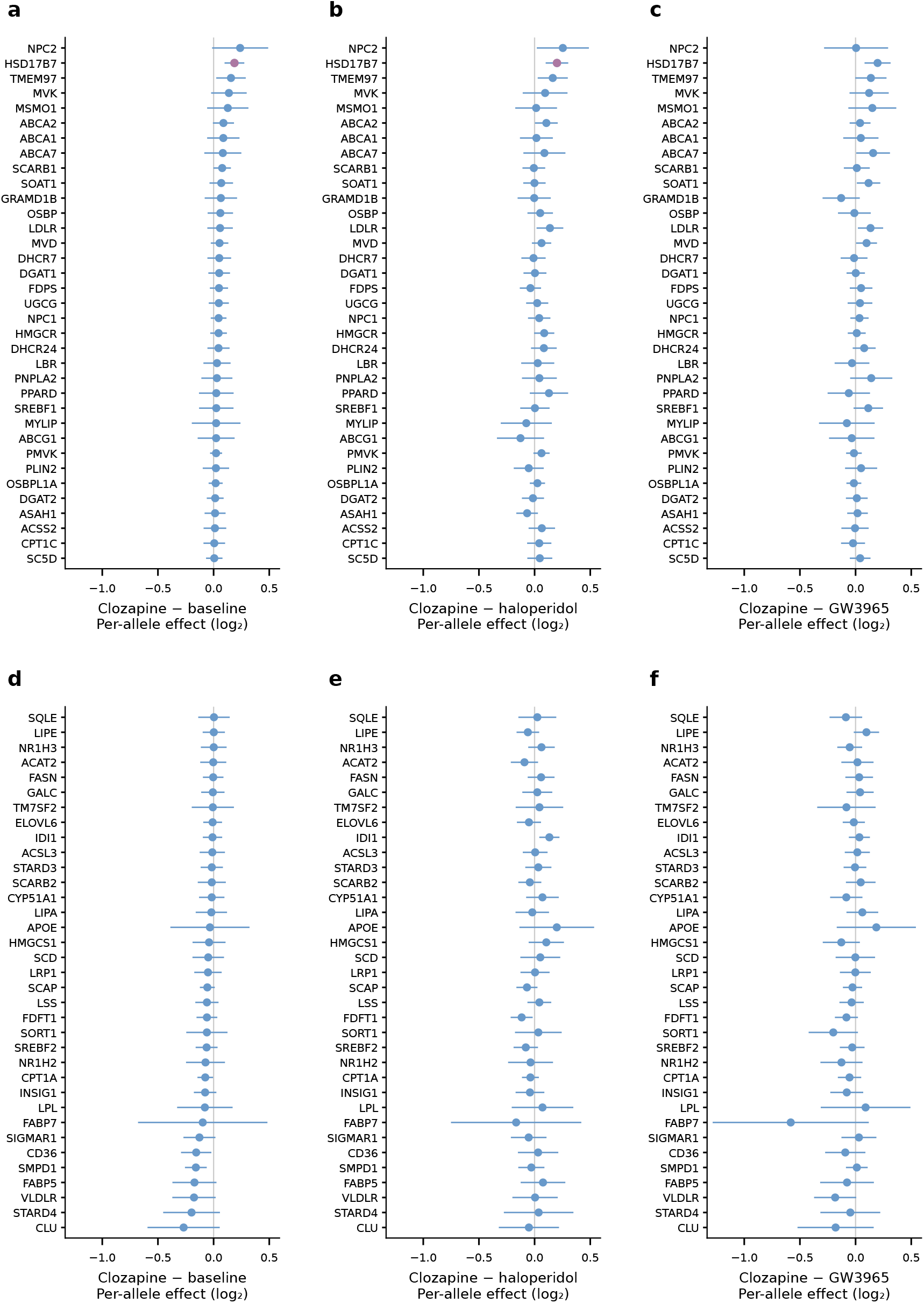
Complete focused lipid-gene response contrasts. **(a–f)** Per-allele effects with 95% confidence intervals for clozapine–baseline **(a,d)**, clozapine–haloperidol **(b,e)** and clozapine– GW3965 **(c,f)** contrasts, comprising 92, 92 and 88 donors, respectively. The first 35 genes appear in (a–c) and the remaining 35 in (d–f), ordered by decreasing clozapine–baseline effect. Axes share limits; purple points indicate FDR < 0.05 within each contrast. Full results are provided in Supplementary Table 10 and source data.

**Extended Data Figure 7.**
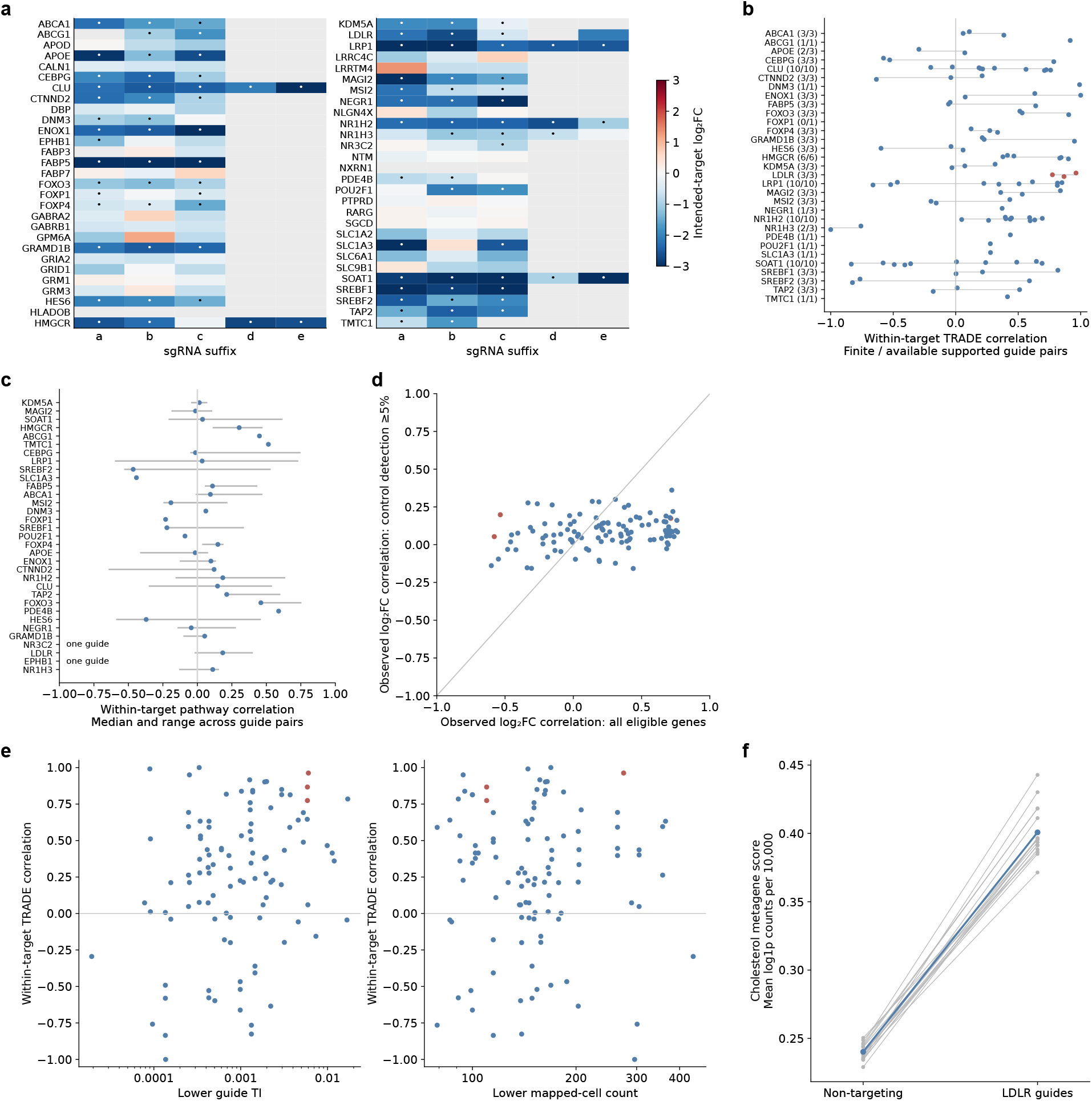
Astrocyte guide support and response concordance. **(a)** Intended-target log_2_ fold-changes across available astrocyte guides. Dots indicate supported repression; grey cells indicate unavailable contrasts. Colors saturate beyond *±*3. **(b)** Within-target TRADE correlations between supported guides. Points represent guide pairs; grey segments span estimates for each target. Labels indicate finite estimates relative to available pairs (105 of 110 overall). **(c)** Within-target agreement of pathway-response profiles. Points and ranges summarize pairwise guide correlations. **(d)** Spearman correlations of gene-level log_2_ fold-changes using all eligible genes versus genes detected in at least 5% of control cells. The diagonal denotes equality. **(e)** Within-target TRADE correlations versus the lower transcriptional impact and lower cell count of each guide pair. *LDLR* pairs are highlighted in red in (b,d,e). **(f)** Cholesterol metagene scores in non-targeting controls and supported *LDLR*-guide cells across 16 sequencing lanes. Scores summarize normalized expression of pathway genes excluding *LDLR*. Grey lines connect measurements within lanes; blue indicates the across-lane mean. Source results are provided in Supplementary Table 11.

**Extended Data Figure 8.**
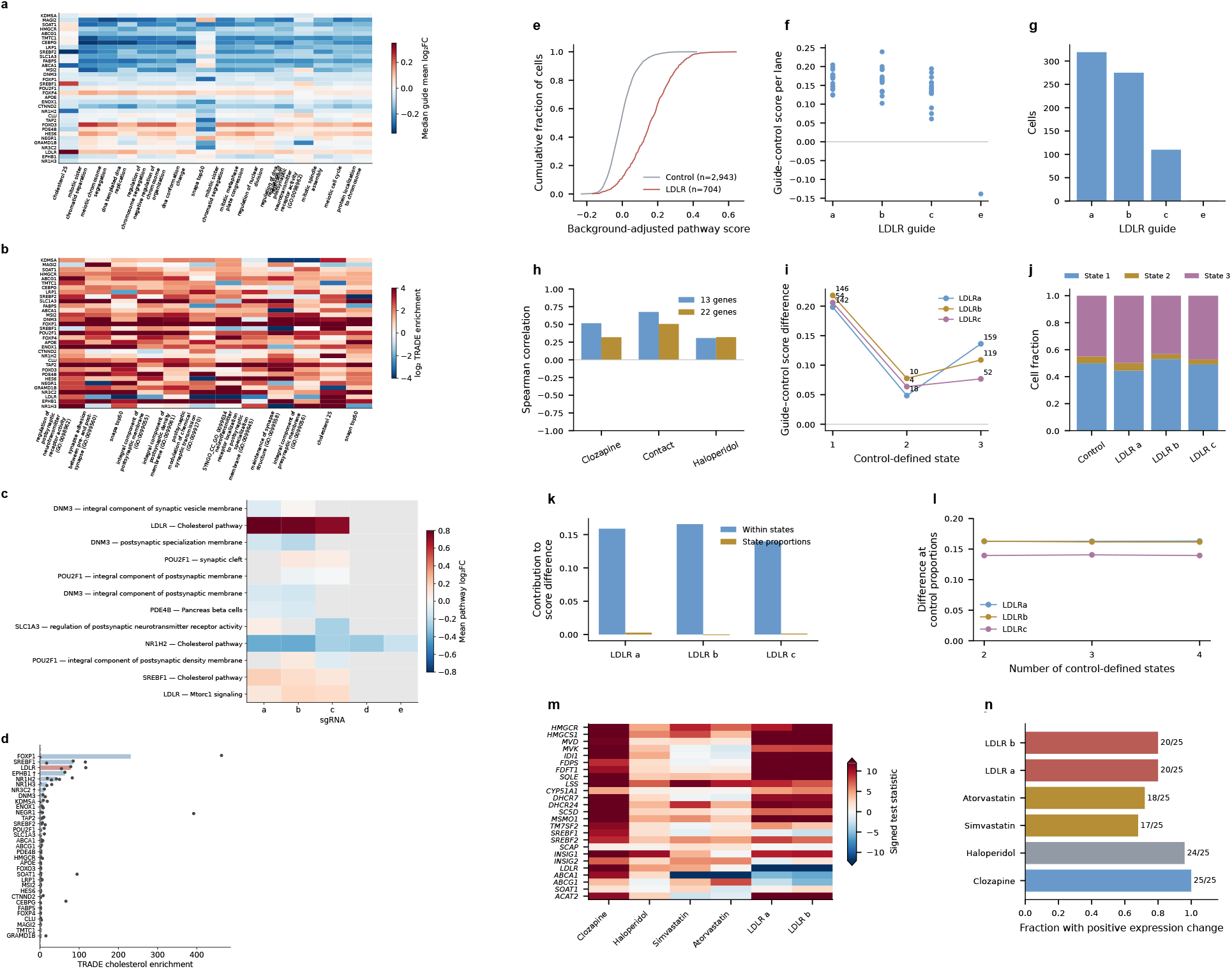
Astrocyte pathway responses and LDLR cell-state and comparator analyses. **(a)** Mean log_2_ fold-changes across selected predefined gene sets and 33 targets with supported repression. **(b)** TRADE concentration estimates across representative SynGO and custom gene sets. Colors show log_2_ enrichment. **(c)** Individual-guide mean log_2_ fold-changes for the target–pathway combinations in Figure 6d. **(d)** Cholesterol-pathway TRADE enrichment across 33 targets. Bars show guide medians; points represent guides. The dashed line marks enrichment of 1. Red highlights *LDLR*; daggers identify single-guide targets. **(e)** Background-adjusted cholesterol-score distributions for 704 cells carrying supported *LDLR* guides and 2,943 non-targeting controls. **(f)** Guide-minus-control mean scores by sequencing lane. **(g)** Cell counts per guide. Scores in (e,f,i–l) summarize expression of 24 cholesterol-pathway genes, excluding *LDLR*, relative to expression-matched background genes. **(h)** Spearman correlations between *LDLR*-repression and contact/drug response statistics using the historical 13-gene set or 22 shared downstream cholesterol genes. **(i)** Mean guide–control score differences within three cell states defined from control expression. Labels indicate guide-cell counts; control counts are 1,469, 148 and 1,326. **(j)** Cell-state proportions among controls and each guide. **(k)** Contributions of within-state expression differences and altered state proportions to the overall cholesterol-score difference. **(l)** Within-state score differences standardized to control proportions across two-, three- and four-state partitions. **(m)** Signed gene-level test statistics across 25 cholesterol-pathway genes under clozapine, haloperidol, simvastatin, atorvastatin and *LDLR* repression. Colors saturate beyond *±*12. **(n)** Fractions of genes with positive response estimates, with counts indicated. Source results are provided in Supplementary Table 11 and accompanying source data.

**Extended Data Figure 9.**
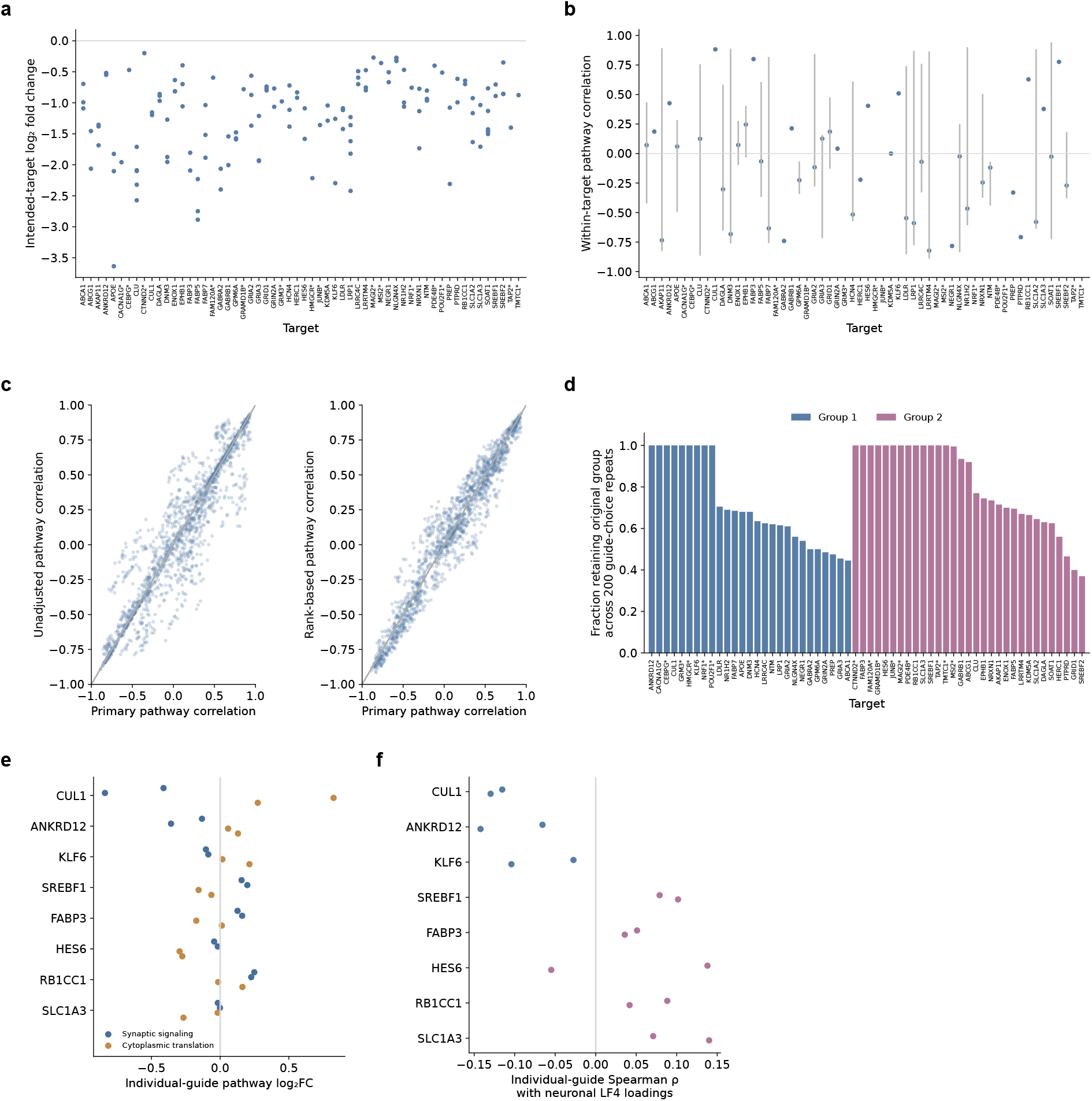
Neuronal guide support and response robustness. **(a)** Intended-target log_2_ fold-changes for 133 supported guides across 58 targets. Asterisks identify single-guide targets. **(b)** Within-target Spearman correlations across 886 pathway-response profiles. Points show medians; lines span correlations between supported guide pairs. **(c)** Comparison of target-pair pathway correlations using the primary profiles versus unadjusted pathway means (left) or expression-adjusted gene ranks (right). Points represent target pairs; diagonals denote equality. **(d)** Fraction of 200 repetitions in which each target retained its original group assignment after selecting one supported guide per target and repeating the clustering. Colors indicate the groups in Figure 7c,d; asterisks mark single-guide targets. **(e)** Individual-guide mean log_2_ fold-changes underlying Figure 7e. Blue indicates synaptic signaling; orange indicates cytoplasmic translation. **(f)** Individual-guide Spearman correlations with neuronal LF4 loadings underlying Figure 7g. Colors indicate the groups in Figure 7c,d. Source estimates are provided in Supplementary Table 12.

**Extended Data Figure 10.**
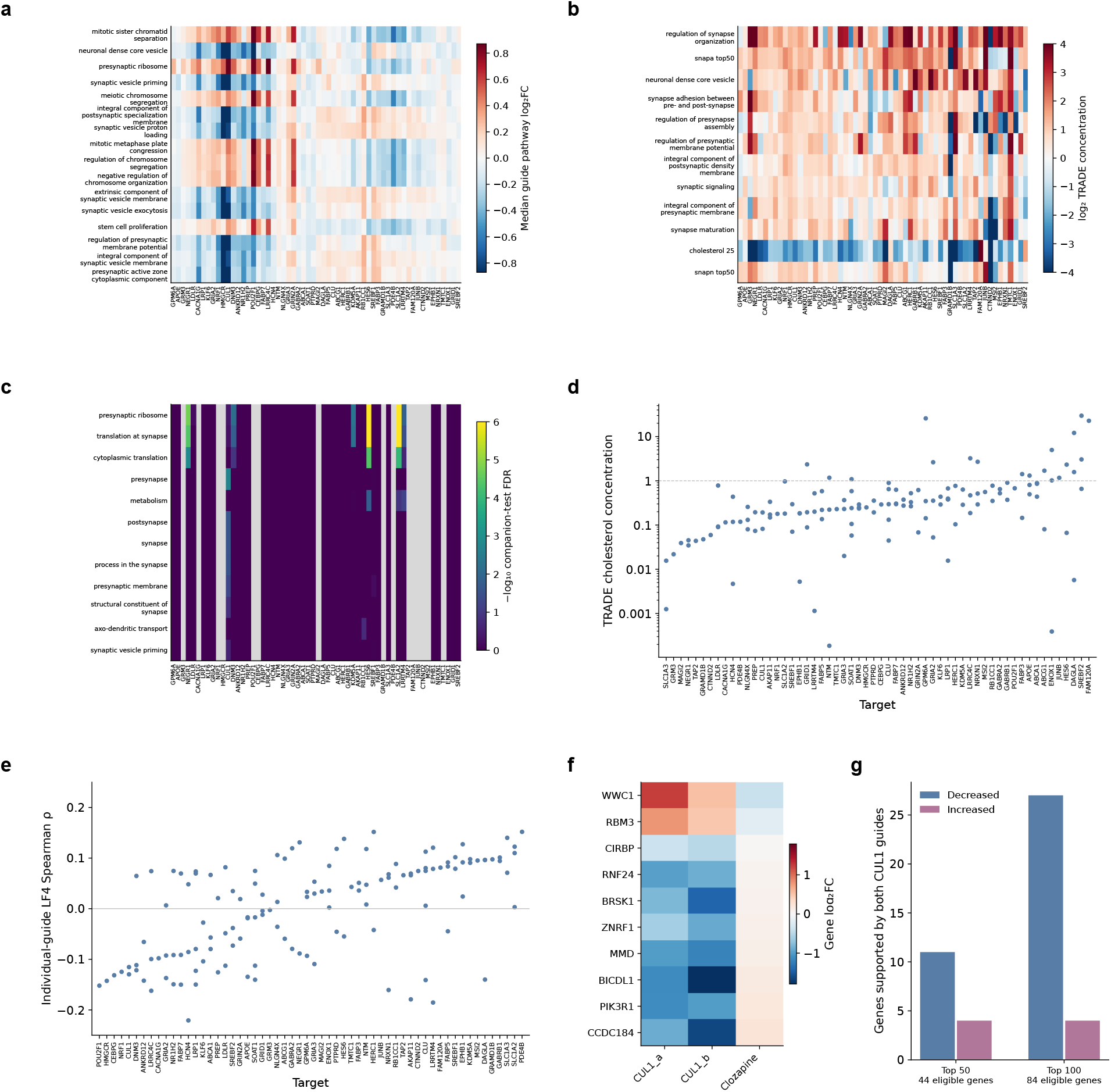
Neuronal pathway concentrations and SNAP correspondence. **(a)** Mean log_2_ fold-changes across 16 representative pathways and 58 supported targets. Values show median guide responses; target ordering follows Figure 7c. **(b)** Median TRADE concentration across selected SynGO, cholesterol and SNAP gene sets. Colors show log_2_ concentration, saturated beyond *±*4. **(c)** Magnitude-enrichment results for the twelve gene sets with the lowest target-level FDR. Colors show *−*log_10_ FDR, capped at 6; grey indicates unavailable multiguide tests. **(d)** Cholesterol-pathway TRADE concentration for supported neuronal guides, ordered by target median. The vertical axis is logarithmic; the reference line marks concentration of 1. **(e)** Individual-guide Spearman correlations between neuronal log_2_ fold-changes and published LF4 loadings, ordered by target median. **(f)** Ten top-100 neuronal LF4 genes with significant opposing expression changes under *CUL1* repression and clozapine. Columns show effects for both *CUL1* guides and clozapine. **(g)** Numbers of top-50 and top-100 neuronal LF4 genes with concordant increases or decreases at adjusted *P* < 0.05 under both *CUL1* guides. The corresponding measured sets contain 44 and 84 genes. Source estimates are provided in Supplementary Table 12.

## Supplementary Tables

**Supplementary Table S1. Village representation and donor composition.** Donor representation during pooled culture, passaging, cryopreservation and differentiation. Worksheets contain Census-seq measurements, single-cell donor-recovery summaries and demographics of the primary pharmacological-screen cohort.

**Supplementary Table S2. Cohort and donor metadata.** Donor and cell-line characteristics for the 44-donor primary pharmacological screen and donor records for the expanded neuronal village. The expanded-village sheet identifies membership in each expression matrix and analysis subset. Recorded demographics and model covariates are listed separately; blank demographic fields indicate unavailable information. Field definitions and sources are included in the workbook.

**Supplementary Table S3. Pharmacological-screen treatment conditions.** Compounds and other perturbations used in the primary screen, with concentrations, exposure durations and experimental annotations.

**Supplementary Table S4. Transcriptional impact and pathway concentration across perturbations.** TRADE estimates of transcriptome-wide impact, associated uncertainty and response dispersion (Me, the effective number of differentially expressed genes) in neurons and astrocytes. Gene-set results report the concentration of transcriptional impact within biological pathways.

**Supplementary Table S5. NeuroPainting experimental design and morphology programs.** Treatment conditions, well metadata, image-derived features and normalized well profiles for the astrocyte NeuroPainting experiment. Worksheets also report compound-associated changes in the eight morphology programs and the corresponding statistical tests.

**Supplementary Table S6. Co-culture latent factors and gene-set enrichment.** Latent-factor scores and treatment summaries from the human-neuron–mouse-astrocyte co-culture experiment, together with the gene-set enrichment results displayed for astrocytic LF_9 and neuronal LF_11.

**Supplementary Table S7. Synaptic-program membership and expression responses.** Gene membership and expression effects linking LF_11-associated synaptic pathways to the published neuronal co-culture program. Worksheets include shared-gene effects and comparisons of neuronal gene-set responses across treatments.

**Supplementary Table S8. Baseline expression and cholesterol-pathway responses.** Associations between baseline pathway or individual-gene expression and subsequent cholesterol-pathway responses, including comparisons between treatments. Worksheets contain split-gene analyses, cross-validated prediction results, and robust, bootstrap and donor-omission analyses of the ACAT2 association.

**Supplementary Table S9. Cis-eQTL associations and external agreement.** Baseline cis-eQTL catalogue, condition-specific associations in matched donors and cis-eGene counts. External comparisons summarize directional agreement with GTEx brain eQTLs and the independent neuronal dataset from Liang et al.

**Supplementary Table S10. Genetic effects on treatment-associated expression changes.** Genotype–response associations for the focused 70-gene lipid-related family, condition-specific genetic effects and the cholesterol-pathway response scan. Sensitivity analyses examine expression transformation, uncertainty estimates and donor influence, including the HSD17B7 association.

**Supplementary Table S11. Astrocyte CRISPRi responses.** Guide-level target repression, transcriptional impact, pathway concentration and response similarity across astrocyte perturbations. Additional worksheets provide LDLR-associated cholesterol-gene responses, single-cell and sequencing-lane summaries, cell-state analyses and comparisons with neuronal contact and drug responses. Companion gene-set tests across correlation settings are included with the table.

**Supplementary Table S12. Neuronal CRISPRi responses.** Guide-level transcriptional impact, supported target repression, pathway-response similarity and TRADE pathway concentration across neuronal perturbations. Worksheets include clustering stability, synaptic and translation-related responses, associations with neuronal LF4 loadings and CUL1–clozapine comparisons. Guide aliases document the corrected target labels. Companion gene-set tests across correlation settings are included with the table.

## Methods

### Human induced pluripotent stem cell cohort and culture

Human induced pluripotent stem cell (iPSC) lines were obtained from the California Institute for Regenerative Medicine (CIRM) hPSC Repository ^45^ or reprogrammed at the Harvard iPS Core from peripheral blood mononuclear cells collected at McLean Hospital by the Deborah Levy laboratory. Cell-line information and donor metadata are provided in Supplementary Table S2. McLean Levy lines are available through the NIMH Repository and Genomics Resource and the Stanley Center Stem Cell Resource at the Broad Institute. The study followed applicable ethical guidelines and institutional review board approvals. iPSCs were maintained in mTeSR1 medium (Stemcell Technologies, 85850) on Geltrex-coated six-well plates (Thermo Fisher, A1413201; VWR, 62406-161), passaged with Accutase (Stemcell Technologies, 07920) and cultured at 37°C in 5% CO_2_.

The primary pharmacological screen included 44 donors. The expanded neuronal screen used a separate pooled village, with overlapping donor subsets for each analysis: 86 donors for treatment-response profiles, 107 for baseline–clozapine associations, 99 for baseline–clozapine–GW3965 comparisons, 92 for baseline–clozapine–haloperidol comparisons and 88 with measurements in all four conditions. Paired genetic analyses included 92 genotyped donors measured at baseline and under both antipsychotics. CRISPRi experiments used WTC11-derived cells.

### Neural progenitor and excitatory neuron differentiation

iPSCs were differentiated into cortical glutamatergic neurons as previously described^17^. On day 0 (D0), cells were plated at 1 *×* 10^6^ cells per well in six-well plates. On D1, medium was replaced with neural induction medium containing 500 mL DMEM/F12 (1:1; Gibco, 11320-033), 5 mL GlutaMAX (Gibco, 35050-061), 7.5 mL 20% dextrose (Sigma, 1181302) and 5 mL N2 supplement (Invitrogen, 17502048), supplemented with 10 µM SB431542 (Stemcell Technologies, 72234), 2 µM XAV939 (Stemcell Technologies, 72672), 100 nM LDN-193189 (Stemcell Technologies, 72147) and 2 µg/mL doxycycline hyclate (Sigma, D9891). Medium was replaced on D2 and D3. Zeocin (1 µg/mL) was applied for 1–2 days during neural induction.

On D4, neurons were co-cultured with mouse glia (TransnetYX, C57EASTWB) to promote maturation and synaptic connectivity^29,46^. Cultures were maintained in medium containing 500 mL Neu-robasal (Gibco, 21103-049), 5 mL GlutaMAX, 7.5 mL 20% dextrose and 2.5 mL MEM non-essential amino acids (Invitrogen, 11140050), supplemented with B27 (50*×* stock; Gibco, 17504-044), BDNF, CNTF and GDNF (10 ng/mL each; R&D Systems, 248-BD/CF, 257-NT/CF and 212-GD/CF) and 2 µg/mL doxycycline hyclate. Floxuridine (10 µg/mL; Sigma-Aldrich, F0503-100MG) was added from D4 to D5. Neural progenitor samples were collected on D4 before addition of mouse glia.

### Astrocyte differentiation

Astrocytes were differentiated as previously described ^18^. On D0, iPSCs were plated at 1 *×* 10^6^ cells per well in six-well plates. On D1, medium was replaced with N2 medium containing 500 mL DMEM/F12 (1:1; Gibco, 11320-033), 5 mL GlutaMAX (Gibco, 35050-061), 7.5 mL 20% dextrose (Sigma, 1181302) and 5 mL N2 supplement B (Stemcell Technologies, 07156), supplemented with 10 µM SB431542, 2 µM XAV939, 100 nM LDN-193189 and 2 µg/mL doxycycline hyclate. Medium was replaced on D2 with the addition of 1 µg/mL zeocin. On D3, neural progenitor-like cells were dissociated with Accutase (Innovative Cell Technology, AT104-500) and replated at 15,000 cells/cm^2^ on Geltrex-coated plates in Astrocyte Medium (ScienCell, 1801) containing 10 µM Y27632. Cells were maintained in Astrocyte Medium for 30 days.

### Drug treatments

Treatment concentrations and exposure durations are listed in Supplementary Table S3. Concentrations differed between the initial screen and the dose and time-course experiments. LXR-623 was excluded from the extended antipsychotic analyses. In the expanded neuronal village, separate wells were treated for 72 h with clozapine (800 ng/mL), haloperidol (1 µM) or GW3965 (50 ng/mL). Untreated wells served as baseline controls. Solvent identity and final concentration were matched across drug treatments. In the neuron–glia co-culture transcriptional experiment, clozapine was tested at 400, 800 and 1,600 ng/mL for 24, 48 and 72 h. The initial screen included 7 h and 24 h IFN-*α* exposures and an inflammatory cocktail containing IL-1*α*, TNF-*α* and C1q.

### Cell villages and single cell RNA sequencing

On D0, equal numbers of iPSCs from each donor (1 *×* 10^6^ cells per donor) were pooled and differentiated into neurons or astrocytes using the protocols above. Cells were harvested at D28 for neurons and D30 for astrocytes. We prepared 60,000 cells for sequencing using 10x Chromium Single Cell 3*^′^* Reagents v3 and sequenced libraries on an Illumina NovaSeq 6000 S2 flow cell with 2 *×* 100-bp paired-end reads.

Reads were processed with the Drop-seq workflow^47^, aligned to GRCh38 and retained at mapping quality *≥*10. Donors were assigned with Dropulation using quality-filtered reference variants. For each droplet, the algorithm calculated donor genotype likelihoods from UMI-supported alleles across transcribed variant sites and assigned the donor with the highest combined likelihood.

### Bulk RNA sequencing

RNA was extracted from D28 neuron–astrocyte co-cultures or D30 astrocytes using the RNeasy kit (Qiagen, 74106). Libraries were prepared by Smart-seq2 and sequenced at the Broad Genomics Platform. Adapter sequences were marked with Picard MarkIlluminaAdapters v3.1.0. Reads were aligned with STAR v2.5.2a to a combined human GRCh38 and mouse GRCm38 reference. Picard MergeBamAlignment was used to merge alignment information, and PCR duplicates were marked. Transcript abundances were quantified with RSEM v1.3.3 using rsem-calculate-expression. Human and mouse count matrices were generated separately for each batch and combined across batches within species.

### Cholesterol and apolipoprotein measurements

Astrocytes from two donor-derived lines were generated in one differentiation per line and treated with clozapine (800 ng/mL) for 72 h. Total cholesterol, including free and esterified cholesterol, was measured in cells and conditioned medium using the fluorescence readout of the Cholesterol/Cholesteryl Ester Assay Kit (Abcam, ab65359). APOE and CLU in conditioned medium were measured by ELISA (Abcam, ab108813 and ab174447). Measurements were background-corrected and divided by the pooled control mean across both lines.

Each measurement came from a separate well. APOE groups contained eight wells, four per line. CLU groups contained six vehicle wells, three per line, and eight wells per drug treatment, four per line. Intracellular cholesterol groups contained six wells, three per line; conditioned-medium cholesterol groups contained four wells, two per line. Treatment and control wells were balanced across plate positions.

### High-density microelectrode array recordings

Neuronal activity was measured using the MaxTwo high-density microelectrode array system (MaxWell Biosystems). Six-well chips were incubated with 1% Terg-a-zyme (Sigma) for 2 h at room temperature, washed three times with deionized water and sterilized with 70% ethanol for 1 h. After three washes with sterile water, wells were primed with 1.2 mL complete neuronal medium for 2 days at 37°C in 5% CO_2_. Chips were then coated with 0.1 mg/mL poly-D-lysine (Sigma) for 3 h at 37°C, washed, dried for 1 h and coated with 50 µL Geltrex for 1 h at 37°C. We plated 300,000 neural progenitors and 60,000 mouse glia in 50 µL medium at the center of each chip. Cultures were maintained at 37°C in 5% CO_2_ with half-medium changes three times per week.

Cultures were treated with clozapine (800 ng/mL) or vehicle for 24 h. Recordings were acquired at 37°C in 5% CO_2_ using the Activity Scan Assay followed by the Network Assay. Each well was measured once. The experiment used one cell line and one differentiation, with treatment and control wells balanced across plate positions. Mean inter-spike interval (ISI) was measured in seven wells per condition; within-burst ISI was measured in five vehicle and seven clozapine wells. Two-sided Welch tests compared treatments, with Holm correction across the two measures.

### Census-seq

DNA was extracted from fluorescence-activated cell-sorted samples using a column-free procedure. Samples were resuspended in 300 µL Cell Lysis Solution (Qiagen, 158906) with 2 µL Proteinase K (New England Biolabs, P8107S) and incubated overnight at 56°C. RNase A (1.5 µL; Qiagen, 158922) was added for 30 min at 37°C. Samples were placed on ice, mixed with 200 µL Protein Precipitation Solution (Qiagen, 158910), vortexed and centrifuged at 13,200 rpm for 10 min at 4°C. Supernatants were transferred to 300 µL ice-cold isopropanol containing 0.5 µL glycogen (Qiagen, 158930) and centrifuged again. DNA pellets were washed with 300 µL 70% ethanol, centrifuged at 14,000 rpm for 5 min at 4°C, dried for 10 min and rehydrated in 15–20 µL water for 1 h at 55°C. Libraries were prepared with TruSeq Nano DNA Library Prep for NeoPrep (Illumina, NP-101-9001DOC) or Nextera DNA Library Prep (Illumina, FC-121-1030). Up to 16 libraries were pooled per run and sequenced on an Illumina NextSeq 500 using a 75-cycle high-output kit to generate single 85-bp reads.

### sgRNA library design and cloning

The sgRNA library targeted 94 genes: 14 cholesterol-biosynthesis enzymes and regulators, 42 genes prioritized by astrocytic SNAP-a cNMF loadings^35^, and 38 genes nominated by common-variant schizophrenia GWAS or rare-variant SCHEMA associations. We selected three to five guides per gene using CRISPick, prioritizing predicted on-target activity and minimizing predicted off-target effects. Non-targeting guides were included as controls. Oligonucleotides were cloned by Golden Gate assembly into a lentiviral vector containing a U6-driven sgRNA cassette, verified by Sanger sequencing and pooled in equimolar amounts. Differential-expression results were available for 86 of the 94 targets across the three profiled cell states.

### Lentivirus production

HEK293T cells were transfected with the pooled sgRNA transfer plasmid, psPAX2 packaging plasmid and pMD2.G envelope plasmid. Viral supernatants were collected at 48 and 72 h, filtered through 0.45-µm membranes and concentrated by ultracentrifugation. Titers were determined by serial-dilution transduction followed by flow cytometry or antibiotic selection.

### CRISPRi cell line and transduction

CRISPRi experiments used WTC11-Ngn2-dCas9-KRAB iPSCs expressing dCas9-KRAB and doxycycline-inducible NGN2. For the neuronal screen, one pooled iPSC culture was transduced with the sgRNA library at a multiplicity of infection of approximately 5, differentiated into excitatory neurons and co-cultured with mouse glia from D4. Cells were collected for single-cell RNA sequencing at D28. For the astrocyte screen, one pooled culture was transduced at D25 at the same multiplicity of infection and collected at D30, after 5 days of repression.

### CRISPRi sequencing and guide assignment

Cells were processed with 10x Chromium Single Cell 3*^′^* Reagents v3 and sequenced on an Illumina NovaSeq 6000. Guide identity was determined using the CROP-seq strategy ^14^, in which a polyadenylated transcript containing the guide sequence is captured alongside cellular mRNAs. UMI-deduplicated guide counts were assigned to cell barcodes within each library. Guide-assignment purity was calculated as the fraction of guide counts contributed by the most abundant sgRNA, with a threshold above 90%. The final analysis retained cells assigned to a single guide and excluded cells with no guide or multiple detected guides. Guide assignments were linked to expression profiles by cell barcode and annotated by intended target or non-targeting control status. The 16 astrocyte sequencing lanes sampled one pooled culture; neuronal libraries likewise sampled one pooled culture.

### Transcriptome wide impact and perturbation similarity

We analyzed 26 condition-by-cell-type differential-expression tables from the primary pharmaco-logical screen. TRADE estimated transcriptome-wide impact (TI) and the effective number of differentially expressed genes (M_e_) from gene-level effects and standard errors. TI summarizes effect variance, whereas M_e_ summarizes its distribution across genes. Gene-set analyses used genes shared across all 26 comparisons with finite effects and standard errors and absolute log_2_ fold-change *≤*10. The MSigDB v2023.1 Gene Ontology biological-process collection was restricted to sets with at least 50 eligible genes, yielding 1,571 sets. Pathway enrichment was calculated as the fraction of TI attributed to a gene set divided by its fraction of measured genes.

We compared TI between cell types for the nine shared conditions and calculated Spearman correlations of gene-level response statistics between perturbations within each cell type. To test similarity within functional categories, we permuted category labels across agents while preserving each agent’s label across cell types. The statistic was the difference between median same-category and different-category correlations. We used 20,000 permutations, random seed 0 and the plus-one correction for empirical P values. Neuronal donor agreement was summarized by the median correlation across donor pairs for each perturbation. Its association with TI was tested across the 14 neuronal perturbations.

### Biochemical assay statistics and antipsychotic expression comparisons

Drug–vehicle differences in biochemical assays were tested by ordinary least squares with treatment and donor line as fixed effects. Line A corresponded to the first half of wells in each assay–treatment group and line B to the second half. Two-sided tests used HC3 standard errors and residual degrees of freedom. Holm correction covered six contrasts: clozapine versus vehicle for cholesterol in cells and medium, APOE and CLU, and haloperidol versus vehicle for APOE and CLU. Measurements were analyzed on their recorded normalized scale without further rescaling. Conventional model intervals and Welch tests were also calculated.

For the extended antipsychotic expression comparison, bulk counts were converted to counts per million (CPM). Two-sided Wilcoxon rank-sum tests compared clozapine with vehicle, with Benjamini– Hochberg correction across the six displayed genes. Comparisons for other compounds used nominal P values. Doses and durations were pooled within treatment.

### Morphological profiling

The NeuroPainting dataset contained 246 wells and 770 image-derived features. We retained 762 features with finite values. Features were normalized within plates by the median and median absolute deviation, followed by an inverse-normal transformation. Positive and negative components were separated to obtain 1,524 nonnegative columns. Non-negative matrix factorization used eight components, NNDSVD initialization, random seed 0, tolerance 10*^−^*^6^ and a maximum of 4,000 iterations. Program activities were adjusted for cell count and categorical exposure duration. Drug–vehicle Wilcoxon rank-sum tests were corrected across 64 program–compound comparisons. Program composition was summarized using the 100 highest-loading features, with channel frequencies compared against the 762 input features. Experimental conditions, measured features and program results are provided in Supplementary Table 5.

Representative images were selected from the 48 h condition. For each treatment, we chose the well nearest its group median program activity and sampled crop positions before visual inspection. Images were displayed using identical intensity limits. Crops spanned 380 pixels at 0.598 µm per pixel, with 50-µm scale bars.

### Comparison with published co-culture signatures

Drug-response results were matched to the Pietiläinen and Vartiäinen co-culture datasets using one-to-one mouse–human orthologues and gene-symbol aliases. Co-culture-induced genes had positive log_2_ fold-change and adjusted P < 0.05 in the source study. We compared the fraction of positive drug effects and mean log_2_ drug effects between induced genes and the remaining measured genes. Analyses were repeated using common gene backgrounds and after excluding cholesterol or broader sterol genes.

Two-sided Fisher exact tests compared positive and nonpositive clozapine effects within and outside each signature, with Holm correction across the two signatures. We also compared signed clozapine Wald statistics using a variance-inflation factor of 1 + (m *−* 1)*ρ*, where m is gene-set size and *ρ* is the within-set correlation parameter ^48^. We evaluated *ρ* at 0, 0.001, 0.005, 0.01 and 0.05. Counts and results for each parameter setting are included in Figure 3 Source Data.

The cholesterol-related and synaptic-associated gene sets comprised 15 and 13 genes from Pietiläinen Supplementary Tables 9 and 12, respectively. The former was enriched among glial genes induced by co-culture; the latter was associated with the neuronal maturation eigengene. Functional categories were selected in order of enrichment significance, requiring at least ten mapped genes and excluding categories sharing more than half their members with a previously retained category. Fourteen categories were displayed. Time-course comparisons used the 48 h astrocyte monoculture as the reference for every co-culture duration. For the additional drug comparisons in Figure 3b,e,f, we used primary-screen differential-expression log_2_ fold changes for clozapine, haloperidol, simvastatin and atorvastatin. In the separate extended astrocyte experiment, counts were normalized to CPM and transformed as log_2_(CPM + 1). For each library, we subtracted the mean vehicle expression at the same exposure duration, averaged replicates within each dose–duration condition, and then averaged equally across conditions. Genes were retained if they reached 1 CPM in at least half of vehicle libraries. These extended-screen summaries were not assigned new P values. Biosynthesis summaries used *HMGCR, HMGCS1, MVD, MVK, IDI1, FDPS, FDFT1, SQLE, LSS, CYP51A1, DHCR7, DHCR24, SC5D, MSMO1, TM7SF2* and *ACAT2* ; transport/export summaries used *ABCA1, ABCG1, APOE* and *CLU*. Panel summaries used genes measured in all four treatments within each experiment and in the Vartiainen reference. The other contact-induced group excluded the published 15-gene cholesterol set. Gene membership and per-gene estimates are included in the accompanying Figure 3 source data.

### Co culture latent factors and neuronal expression

The PEER analysis generated 15 factors. Libraries were grouped by treatment, dose and duration, and factor scores were summarized by the median within each condition. The heatmap included 78 conditions across nine treatments, with scores standardized within each factor. Treatment summaries show means and standard errors across condition-level scores.

For LF_9 and LF_11, we displayed the ten gene sets with the highest positive normalized enrichment scores. LF_11 leading-edge genes were compared with the published neuronal co-culture program among 1,442 SynGO-annotated genes measured in both datasets. We calculated overlap for each term and for the non-redundant union of leading-edge genes.

Human-neuron expression in the neuron–mouse-glia experiment was analyzed using edgeR filtering and TMM normalization followed by limma-voom. Models included treatment and categorical exposure duration, pooling doses and using DMSO as the reference. Untreated samples were excluded. We retained moderated effects and standard errors. Expression plots used log_2_(CPM + 1) from libraries with at least one million reads, comprising 26 vehicle and 20 clozapine libraries. Five shared genes were selected to illustrate the observed treatment differences. Results for all shared genes are provided in Supplementary Table 7 and Extended Data Figure 2i–l.

### Cholesterol pathway and donor inclusion

The 25-gene cholesterol-pathway set comprised *HMGCR*, *HMGCS1*, *MVD*, *MVK*, *IDI1*, *FDPS*, *FDFT1*, *SQLE*, *LSS*, *CYP51A1*, *DHCR7*, *DHCR24*, *SC5D*, *MSMO1*, *TM7SF2*, *SREBF1*, *SREBF2*, *SCAP*, *INSIG1*, *INSIG2*, *LDLR*, *ABCA1*, *ABCG1*, *SOAT1* and *ACAT2*. This set includes biosynthetic enzymes, regulators and transport genes. In CRISPRi pathway summaries, the intended target was excluded from its own gene set; *LDLR* scores therefore used 24 genes.

Donor response profiles used treatment coefficients relative to untreated baseline for 86 donors. Gene-level means and 95% intervals were calculated from 5,000 donor-bootstrap resamples, using the same sampled donors across genes and compounds (random seed 42). Baseline-expression analyses used overlapping subsets from the same village: 107 baseline–clozapine pairs, 99 baseline– clozapine–GW3965 triplets, 92 baseline–clozapine–haloperidol triplets and 88 donors measured in all four conditions.

Expression matrices were transformed as log_2_(x + 1) before baseline-expression and paired-response analyses.

### Baseline expression and response associations

We randomly divided the 25 pathway genes into 12 predictor genes and 13 outcome genes 200 times (random seed 42). For each partition, we calculated Pearson correlations between mean baseline expression in the predictor set and mean treatment-induced change in the complementary set. The displayed partition had the correlation closest to the mean across partitions. Comparisons across compounds used the same 88 donors and gene partitions. We summarized correlations by their mean and 2.5th–97.5th percentiles across partitions.

For each pathway gene, an ordinary least-squares model related its baseline expression to the mean clozapine-minus-comparator expression difference across the other 24 genes, adjusting for their mean baseline expression. Both comparator analyses used the same 88 donors. Two-sided tests used n *−* 3 residual degrees of freedom. Benjamini–Hochberg correction was applied across 25 genes per comparator and, separately, across the pooled 50 tests. HC3 standard errors were evaluated using the same degrees of freedom and correction families. Leave-one-donor-out analyses refitted all 25 models and repeated correction. *ACAT2* bootstrap intervals used 10,000 donor resamples (random seed 20260923).

### Prediction and permutation analysis

We compared a ridge-regression model using mean baseline pathway expression with a model that also included all 25 baseline genes after regression on the pathway mean. Prediction was evaluated across 20 repetitions of donor-level five-fold cross-validation (random seeds 0–19). Residualization and scaling were fitted within each outer training fold and applied to held-out donors. Ridge penalties were selected from 30 logarithmically spaced values between 0.001 and 10,000 using analytic leave-one-out error within the training data, with preprocessing held fixed during penalty selection.

Held-out predictions were pooled within each repetition. R^2^ was calculated as one minus the residual sum of squares divided by the total sum of squares around the observed outcome mean. We averaged R^2^ and the paired gain from adding individual-gene expression across repetitions. The 2.5th–97.5th percentiles summarize variation across fold assignments.

For the residual-permutation test, we regressed the outcome on baseline pathway mean, permuted the residuals and added them to the reduced-model fitted values. Both prediction procedures, including penalty selection, were repeated for 999 permutations. We also evaluated a global outcome-shuffle test.

### Expression quantitative trait loci

Cis-eQTL analyses tested variants within *±*100 kb using inverse-normal-transformed expression, donor covariates and 15 expression principal components. Variant-level statistics were combined using ACAT, followed by gene-level Benjamini–Hochberg correction. The baseline catalogue included 13,526 genes. For comparisons across conditions, correction was applied to gene-level ACAT P values for the same 13,331 genes in 92 donors measured at baseline and under both antipsychotics.

Baseline associations were matched by gene and variant to GTEx v8 brain eQTLs and the Liang neuronal dataset. The GTEx comparison comprised 1,610 gene–variant–tissue records across 556 genes; the Liang comparison comprised 1,952 records across 472 gene–variant pairs in nine neuronal contexts. Directional agreement was summarized across matched records. Confidence intervals used 10,000 gene-clustered bootstrap resamples, retaining all contexts for each sampled gene.

Paired genetic analyses tested baseline lead variants for 70 lipid-related genes. Outcomes were clozapine-minus-baseline, clozapine-minus-haloperidol and clozapine-minus-GW3965 differences in log_2_(x + 1) expression. Models included alternate-allele dosage and nine donor covariates representing age, sex, diagnosis, 22q status, source indicators and three genotype principal components. Expression principal components were omitted. The three contrasts included 92, 92 and 88 donors, respectively. Conventional and HC3 confidence intervals used t distributions with residual degrees of freedom. Benjamini–Hochberg correction was applied across the 70 genes separately for each contrast and standard-error specification.

We repeated the paired analyses after omitting each donor, excluding alternate homozygotes and changing the log-transformation pseudocount. Baseline LSS and LDLR plots used catalogue lead variants; paired response models used leads from the matched-donor baseline scan. The catalogue and matched-scan leads differed for LDLR and TMEM97. For the pathway analysis, the mean covariate-adjusted clozapine–baseline difference across 25 genes was tested against 13,066 lead variants in 92 donors, with Benjamini–Hochberg correction across variants.

### CRISPRi guide validation

Differential-expression results comprised 203 astrocyte and 269 neuronal guide contrasts. Target labels NXRN1 and HLADOB were mapped to *NRXN1* and *HLA-DOB* before matching to expression results. Finite intended-target P values were available for 203 astrocyte and 266 neuronal contrasts. Two-sided target P values were Benjamini–Hochberg corrected within cell type. Supported repression required a negative target effect and FDR < 0.05. *IFITM3*, *CASP3*, *CASP7*, *AGTRAP* and *SMARCB1* were excluded after correction because their results are reported separately. The retained supported sets contained 97 guides targeting 33 genes in astrocytes and 133 guides targeting 58 genes in neurons. We also compared guide inclusion using transcriptome-adjusted P values, nominal P values and effect direction.

Astrocyte guide effects shared 29,269 genes with finite estimates and statistics. Target-level pathway summaries weighted supported guides equally and excluded the intended target from downstream gene sets. Gene-level intervals were calculated as the effect *±*1.96 standard errors. Target means and standard errors were calculated across guide-level pathway averages. Original gene-wise adjusted P values were retained.

### CRISPRi transcriptome wide impact and pathway analyses

Screen-wide analyses included all 428 in-scope guide contrasts: 181 astrocyte and 247 neuronal contrasts. Within each cell type, genes were required to have finite log_2_ fold-changes and P values, positive finite standard errors, absolute log_2_ fold-change *≤*10 in every contrast and median base-Mean *≥*1 across contrasts. These criteria yielded 12,520 astrocyte and 11,965 neuronal genes. Each guide’s intended target was then removed from its background and gene sets; other screen targets were retained.

TRADE^20^ was fitted to guide-level effects and standard errors to estimate TI and M_e_. Pathway and complement effect distributions were fitted separately. The pathway variance fraction was calculated as n_s_v_s_/(n_s_v_s_ + n_c_v_c_), where n is the number of genes and v the fitted variance in the set (s) or complement (c). TRADE enrichment was this fraction divided by the pathway’s fraction of background genes. We report point estimates of concentration.

The gene-set collection contained 1,448 GO biological-process sets with at least 50 eligible genes in both cell types, 108 SynGO v1.3 sets with at least 11 eligible genes before target exclusion ^49^, and three cholesterol/SNAP sets. SynGO annotations were propagated through the ontology, identical gene sets were collapsed and no upper size limit was applied. Sets retained at least ten genes after target exclusion. TRADE concentration was calculated for the 108 SynGO sets, three cholesterol/SNAP sets and 50 Hallmark sets. Competitive tests used the 1,559-set GO BP, SynGO and cholesterol/SNAP collection, with Hallmark analyzed separately.

Competitive enrichment was tested with limma cameraPR ^48^, using use.ranks = TRUE. Magnitude tests used absolute log_2_ fold-change divided by its standard error and directional = FALSE; directional tests used signed statistics. The within-set correlation parameter was set to 0.01, with additional analyses at 0.05 and 0.10. Benjamini–Hochberg correction included all guide–set tests within each cell type, test type and correlation setting. For targets with at least two supported guides, the conjunction P value was the maximum across supported guides; directional disagreement was assigned P = 1. Target-level correction included all eligible target–set combinations within each cell type and test setting. Values above one because of numerical rounding were capped at one before correction.

For the TRADE concentration overview, target–set pairs with at least two supported guides were ranked by their minimum guide concentration across the 108 SynGO, 50 Hallmark and three cholesterol/SNAP sets. Within each target, lower-ranked sets were removed when overlap exceeded 50% of the smaller measured set. The top 12 remaining pairs were displayed using guide medians and ranges. Companion plots show each guide’s mean log_2_ fold-change across the same genes. To examine whether individual genes dominated selected responses, we calculated each gene’s share of the summed squared log_2_ fold-changes within its set.

Within-target bivariate TRADE models used 12,469 astrocyte and 11,906 neuronal genes, excluding all screen targets. Models used the combined covariance-mixture specification and estimated sampling covariance. We also calculated Spearman correlations of observed gene-level fold-changes on the same backgrounds. Expression-coverage checks compared observed correlations across all eligible genes with correlations restricted to genes detected in at least 5% of control cells.

### Single cell cholesterol scores and expression states

Counts were normalized to 10,000 per cell and log1p transformed. Cholesterol metagene scores were calculated as the mean expression of the 24 pathway genes excluding *LDLR*. The analysis included 2,943 non-targeting cells and 704 cells carrying *LDLR* guides a, b or c (319, 275 and 110 cells, respectively). Lane-level means were calculated for each guide and control group; the overall summary weighted the 16 lanes equally.

For background-adjusted scores, we subtracted the mean expression of 1,200 expression-matched genes. Matching used 25 rank-based bins defined in control cells and 50 background genes per pathway-gene bin contribution (random seed 20260922). Control comparisons were repeated using pooled controls, equally weighted control guides and omission of individual control guides. The unsupported *LDLR* guide e contributed one cell to the cell-count display.

Cell states were identified in control cells using the 1,000 highest-variance genes with mean log-normalized expression >0.05, excluding pathway genes, score-background genes and screen targets. Features were standardized, projected onto ten principal components and clustered with k-means. Models were fitted to controls and applied to guide-bearing cells. K-means used 20 initializations, random seed 23 and two, three or four clusters. For each guide, the total score difference was decomposed into guide-weighted within-state mean differences and differences attributable to state proportions, using control state means. Within-state differences were also standardized to control state proportions.

### Cross-experiment cholesterol comparisons

Cholesterol-response correlations used signed gene-level statistics for 22 shared downstream genes across *LDLR* guides and contact/drug comparisons. The contact comparison was mouse astrocyte co-culture versus monoculture at 48 h; drug and CRISPRi comparisons used human astrocytes. Genes in this matrix were matched by symbol. We compared results for the mean *LDLR* a/b statistics using the historical 13-gene set and the 22 shared genes. Statin-response heatmaps included all 25 pathway genes, including *LDLR*.

### Neuronal gene level responses and enrichment

Gene-level guide agreement, conjunction tests and Reactome/Hallmark analyses used 8,130 neuronal genes with finite effects and adjusted P values across retained guides and median baseMean *≥*1. All screen targets were excluded. Guide agreement was summarized by Spearman correlation and same-direction gene-level evidence. For count-based analyses, genes required adjusted P < 0.05 in at least two guides and no significant opposite-direction guide.

For *CUL1*, *SREBF1* and *FABP3*, the gene-level conjunction P value was the larger of the two guide P values when effect signs agreed, and one otherwise. Benjamini–Hochberg correction was applied across 8,130 genes separately for each target. The plotted effect was the mean of the two guide effects.

Ranked GSEA used GSEApy v1.1.11 with signed Wald statistics, weight 1, 10,000 gene-set permutations and random seed 20260923. The 754 eligible Reactome and Hallmark v2023.1 sets contained 15–500 genes from the common background and were tested together per guide. Displayed terms required concordant normalized enrichment-score signs and FDR < 0.05 in both guides. Up to two terms per direction were selected by the larger guide FDR, with ties resolved by canonical term and enrichment-score strength. Terms with Jaccard overlap *≥*0.5 or overlap greater than or equal to 70% of the smaller set were removed from the display. Zero empirical FDR values were retained in tables; *−*log_10_ values were capped at four for plotting.

### Schizophrenia gene comparisons

*SV2A* and *ZMYM2* responses were assessed using guide-specific gene-level models. Risk-gene counts used the concordant two-guide criterion defined above. Common-variant genes were defined by genome-wide-significant MAGMA associations and rare-variant genes by SCHEMA q < 0.05. We compared response counts with 10,000 random gene sets matched within expression deciles, additionally matching gene-length and SNP-count tertiles for common-variant genes. All screen targets were excluded from these comparisons.

### SNAP loading and clozapine comparisons

Published multicellular LF4 loadings and cell-specific cNMF SNAP-a and SNAP-n loadings were analyzed separately ^35^. Genes were ranked by their published loadings before intersection with measured genes. Top-30, top-50 and top-100 LF4 lists were nested; unmeasured genes were not replaced. Analyses included all supported guides and targets. The common background required finite effects, positive finite standard errors and median baseMean *≥*1 across available contrasts, excluding all screen targets. This yielded 12,469 astrocyte and 11,906 neuronal genes. Target-level summaries used guide medians. Gene-level agreement required adjusted P < 0.05 and the same effect direction in every supported guide, with at least two guides per target.

We compared gene-level effects with published loadings using Spearman correlations of log_2_ fold-changes and standardized effects (log_2_ fold-change divided by its standard error). Pearson correlations and cosine similarities were also calculated. Analyses were repeated after excluding the 25 cholesterol-pathway genes. Astrocyte and neuronal loadings were matched to their respective cell types.

Clozapine effects were matched to CRISPRi results by cell type. Genes were required to belong to the common background and have clozapine baseMean >5, finite effects and positive finite standard errors. Effect ranks were compared across the background and within the top-50 and top-100 LF4 sets. Standardized drug effects were calculated from log_2_ fold-changes and standard errors. Opposing responses required adjusted P < 0.05 under clozapine and every supported guide, with all guide effects opposite in sign to clozapine.

## Code and data availability

Data is available on NIMH AnVIL repository under the project: Genetic Neuroscience: How Human Genes and Alleles Shape Neuronal Phenotypes with the dbGaP Study Accession: phs002032.v2.p. Software and core computational analysis to align and process sequencing reads and perform donor assignment are freely available at GitHub (https://github.com/broadinstitute/Drop-seq). Published or publicly available software, tools, algorithms and packages are cited with their version numbers in the text and Reporting Summary. Other custom code is available on request from the corresponding authors.

## Author Contributions

Conceptualization: M.T., S.M., and R.N. Investigation and experimental work: M.T., H.X., V.N., D.L., E.P., and D.H. Methodology and analysis: M.T., N.P., A.N., J.N., S.B. Project management: M.H., A.M., and K.I. Supervision: R.N. and S.M. Writing and editing: M.T., R.N., and S.M.

